# Temporal Changes Guided by Mesenchymal Stem Cells on a 3D Microgel Platform Enhances Angiogenesis In Vivo at a Low-Cell Dose

**DOI:** 10.1101/2020.05.16.086298

**Authors:** Dilip Thomas, Grazia Marsico, Isma Liza Mohd Isa, Arun Thirumaran, Xizhe Chen, Bart Lukasz, Gianluca Fontana, Brian Rodriguez, Martina Marchetti-Deschmann, Timothy O’Brien, Abhay Pandit

## Abstract

Therapeutic factors secreted by mesenchymal stem cells (MSCs) promote angiogenesis *in vivo*. However, delivery of MSCs in the absence of a cytoprotective environment offers limited efficacy due to low cell retention, poor graft survival and the non-maintenance of a physiologically relevant dose of growth factors at the injury site. The delivery of stem cells on an extracellular matrix (ECM)-based platform alters cell behaviour including migration, proliferation and paracrine activity, which are essential for angiogenesis. We demonstrate the biophysical and biochemical effects of pre-conditioning human MSCs for 96 hours on a three-dimensional ECM-based microgel platform. By altering the macromolecular concentration surrounding cells in the microgels, the pro-angiogenic phenotype of hMSCs can be tuned in a controlled manner through cell-driven changes in extracellular stiffness and ‘outside-in’ integrin signaling. The microgels tested at a low-cell dose (5×10^4^ cells) in a pre-clinical hindlimb ischemia model showed accelerated formation of new blood vessels with a reduced inflammatory response impeding progression of tissue damage. Molecular analysis revealed that several key mediators of angiogenesis were upregulated in the low-cell dose microgel group, providing a mechanistic insight of pathways modulated *in vivo*. Our research adds to current knowledge in cell encapsulation strategies by highlighting the importance of preconditioning or priming the capacity of biomaterials through cell-material interactions. Obtaining therapeutic efficacy at a low-cell dose in the microgel platform is a promising clinical route that would aid faster tissue repair and reperfusion in ‘no-option’ patients suffering from peripheral arterial diseases such as Critical Limb Ischemia (CLI).

## Introduction

Mesenchymal stem cell (MSC) based therapy is considered as a promising alternative to the administration of growth factors or genes for inducing therapeutic angiogenesis. In the literature, a paracrine-centric mechanism via release of cytokines, chemokines and growth factors by MSCs and its role in high cell survival (1), recruitment of endothelial progenitors (2) and angiogenesis (3, 4) has been demonstrated (5). Several tissue engineering technologies have exploited the paracrine properties of MSCs via cell immobilisation strategies, with the aim of using cells as local drug delivery depots to secrete beneficial factors to induce reparative angiogenesis (6). However, little is currently known about the stem cell phenotype and behaviour as a result of *in vitro* pre-conditioning in artificially engineered extracellular matrices. The local environment where the cells are immobilised often provides essential cues for cell growth and function (7, 8). Extracellular molecules or mimics used as cell immobilisation matrices enhance cell survival and direct differentiation guided by embedded spatio-temporal cues (9). In the context of replicating the native cellular microenvironment, cells experience their microenvironment quite distinctly in three-dimensional (3-D) scaffolds compared to two-dimensional (2-D) scaffolds (10). Cells grown in 3-D matrices exhibit altered phenotypes due to the inhibition of their rapid proliferative behaviour and contact-dependent suppressive mechanisms observed in 2-D. Furthermore, cell morphology and cytoskeletal organisation are important predictors of cell fate and lineage(11). Previous studies have shown that, varying the macromolecular density influence the morphological behaviour of mesenchymal stem cells (12)-(13, 14) in inducing an angiogenic behaviour. These changes in morphology is dependent on the pliability (or stiffness) of the material. A more labile matrix permits tractions in a stiffness dependent manner (15). Cellular interactions in a 3-D environment mediated via integrin ligands and growth factor signaling molecules leads to activation of intracellular signaling pathways and changes in gene regulation and expression (16, 17). Thus, most recent efforts have focussed on engineering 3-D hydrogel-based scaffolds composed of extracellular components to direct stem cell behaviour, which is attributable solely to the inherent bioinstructive properties of extracellular molecules (18). However, the influence of macromolecular density in the fourth-dimension (4-D) on modulating stem cell behaviour has received less attention. Type-I collagen hydrogels can recapitulate the extracellular matrix composition characteristic of several tissues. Moreover, changes in the macromolecular density of collagen may affect matrix rigidity, alter the intracellular organization and modulate integrin expression. Because of the reciprocity between cells and the surrounding microenvironment, collagen macromolecular concentrations are likely to direct cells towards unique phenotypes. The collagen hydrogel-based cell embedding platform tested here might serve as an ideal model to investigate the impact of mechanosensing and integrin expression on the alteration of MSCs property.

This study expands on our previous assessment of cell-matrix interactions in microgel systems(19), and our understanding of the influence of macromolecular concentration on the ‘angiocrine’-angiogenic paracrine phenotype of hMSCs (**Fig. 1**). Furthermore, this effect is tested as an alternative cell-based therapeutic in inducing functional angiogenesis at a cell-dose that is twenty-fold lower than the standard dose in a pre-clinical rodent model of Critical Limb Ischemia (CLI). Cell dose is a key parameter for the clinical success of cell therapy, particularly in the case of bone marrow derived multipotent stem cell treatment for ‘no-option’ CLI patients. A significant reduction in cell-dose mitigates concerns regarding safety and reduces the costs for treating a large number of patients in the clinic. The overall hypothesis of the study was that, *in vivo* therapeutic angiogenesis can be achieved with the delivery of hMSCs at a low-cell dose on a 3-D type-I collagen microgel, used as a platform to prime hMSCs for a proangiogenic response. The specific objectives of the study were to investigate the degree of stiffness of the extracellular environment over a 96-hour pre-conditioning period. We also monitored changes in cell morphology and integrin expression because of differences in macromolecular concentration, and finally, we tested the efficacy of hMSC primed microgels *in vivo* to promote therapeutic angiogenesis in a severe rodent model of CLI.

**Figure 1.**
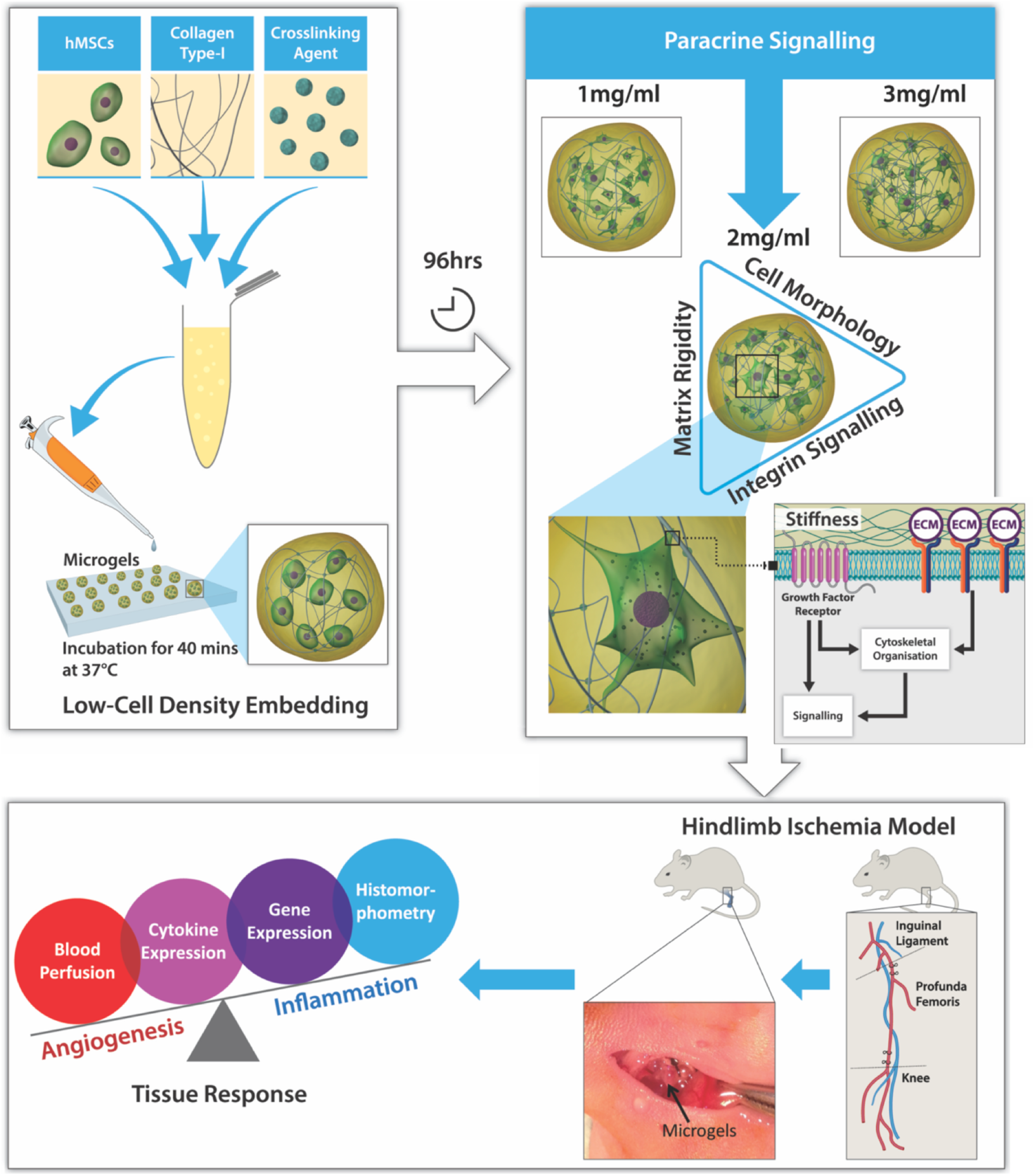
Schematic study design. 3D Microgels embedded with hMSCs at low-cell density were fabricated using microdispensing technique. Pre-conditioning microgels in vitro for 96 hrs modulated the paracrine release of hMSCs dependent on the macromolecular concentration. Further investigation into the integrin expression revealed a pro-angiogenic phenotype of hMSCs embedded in 2mg ml-1 microgels. In vivo testing of 2mg ml-1 microgels in a hindlimb ischemia model showed high angiogenic activity and low inflammatory tissue response contributed to revascularisation and reperfusion.

## Results

### Microgel Fabrication and Experimental Design

Microgels in the form of spherical beads were fabricated by depositing a forming gel solution on a hydrophobic Teflon™ tape as previously described (19, 20). The forming gel solution composed of bone-marrow derived hMSCs, type-I atelocollagen, and poly(ethylene glycol) ether tetrasuccinimidyl glutarate (4S-StarPEG). Microgels of varying collagen concentrations (1mg ml^-1^, 2mg ml^-1^ and 3mg ml^-1^) were fabricated with a cell seeding density of 8 ×10^5^ cells ml^-1^ of the final gel volume. The effect of collagen concentration was investigated using a study design **(Fig. 1)** to understand cell behaviour and matrix remodelling effects due to *in vitro* pre-conditioning.

### Extracellular Microenvironment Drives Substrate Stiffness

Embedding of hMSCs had a significant effect on the remodelling of the extracellular matrix over 96 hours in culture, as evidenced by immunostaining of the whole microgels (**Fig. 2**). The changes in extracellular stiffness were correlated with the localisation of the transcription factor YAP/TAZ during the pre-conditioning period (**Fig. 2a**). Nuclear localisation of YAP/TAZ is indicative of increasing stiffness experienced by the cells, whereas cytoplasmic localisation is observed when cells are on a pliable matrix. A significant difference was observed in the cell morphology and expression of YAP/TAZ cytoplasmic-nuclear translocation between 24 and 96 hours in microgels of different collagen concentrations (**Fig. 2b, c**). A reduced cytoplasmic co-localisation was observed at 96 hours; whereas the nuclear co-localisation increased significantly in 1mg ml^-1^ and 3mg ml^-1^ microgels (p<0.05). No significant changes were observed in 2mg ml^-1^ microgels embedded with hMSCs. Furthermore, changes in the matrix modulus were verified by AFM analysis which showed a greater than a ten-fold increase in microgel stiffness in both 1mg ml^-1^ (1,113Pa) and 3mg ml^-1^(1,007Pa) microgels at 96 hours (**Fig. 2d**). A difference of less than seven-fold was observed in 2mg ml^-1^(452Pa) microgels at 96 hours. In order to understand the influence of gel contraction on YAP/TAZ localisation during the preconditioning period, a myosin blocker blebbistatin was added to the microgels (**SI Appendix, SI Fig. S3a-c**). In all microgel environments, blebbistatin restricted cell spreading compared to controls (**SI Appendix, SI Fig. S3a).** Changes in nuclear localisation in microgel concentrations 1 and 3mg ml^-1^ was not found to be different than controls without blebbistatin. Whereas there was an increase in nuclear YAP/TAZ localisation in 2 mg ml^-1^ after blebbistatin treatment.

**Figure 2.**
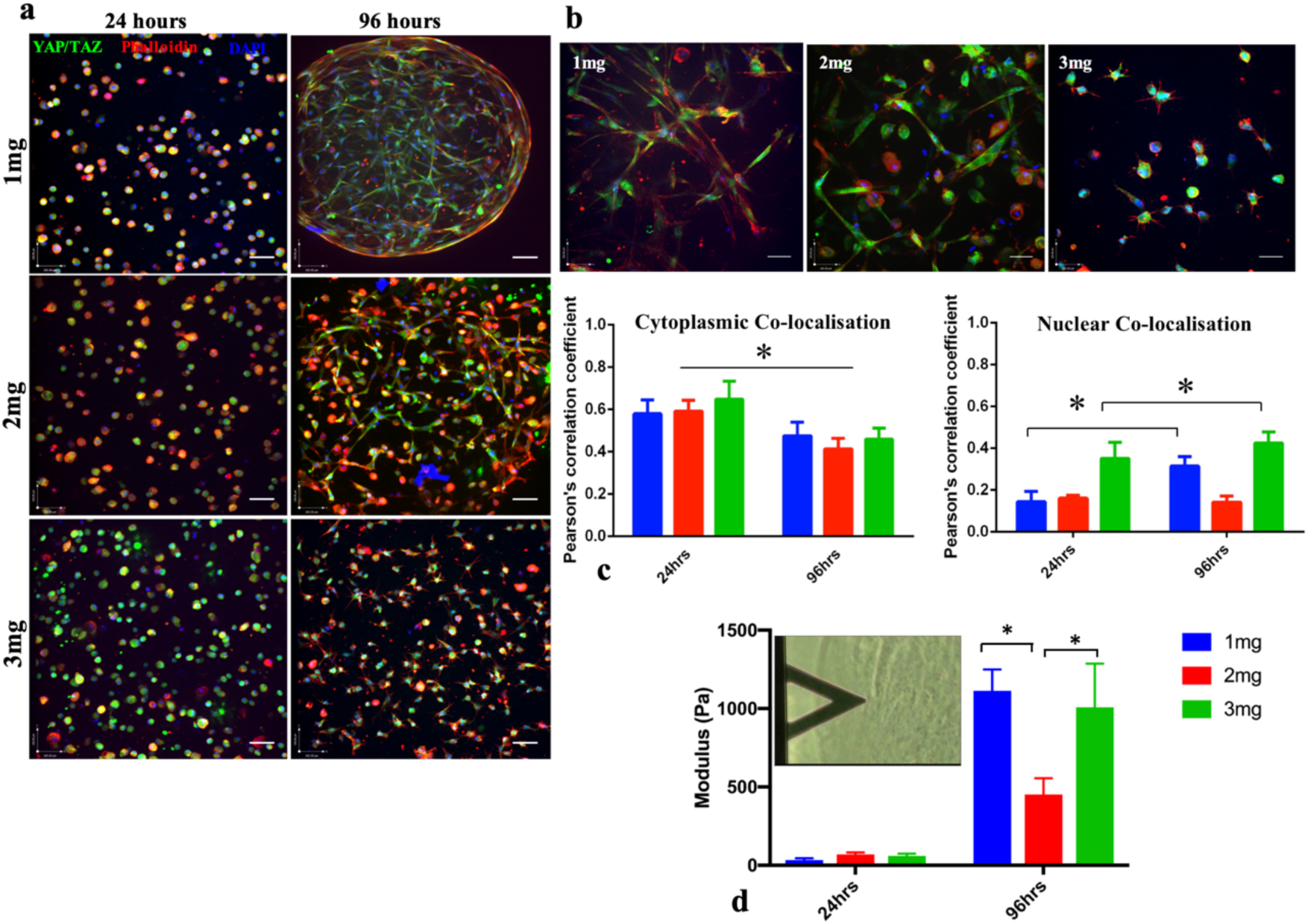
Microgel macromolecular concentration alters the elastic modulus of the cell embedded microgels and localisation of YAP/TAZ. (a) Immunostaining of YAP/TAZ (green), cytoskeleton (red), and nucleus (blue) of human mesenchymal stem cells in 1, 2 and 3mg ml^-1^ microgels at 24 hrs and 96 hrs. Magnification 20X, Scale bar, 100µm. (b) Changes observed in cell morphology and YAP/TAZ (green) localisation, cytoskeleton (red), and nucleus (blue) of human mesenchymal stem cells in 1,2 and 3mg ml^-1^ microgels at 96 hrs. Magnification 40X, Scale bar,100µm. (c) Quantification of cytoplasmic or nuclear YAP/TAZ co-localised with the cytoplasm (Phalloidin) or nucleus (DAPI) using Pearson’s coefficient for co-localisation. Significant increase in nuclear YAP/TAZ co-localisation observed in 1 and 3mg ml^-1^ (n=4). (d) Elastic moduli of hMSC seeded microgels (1, 2, and 3mgml^-1^) measures via atomic force microscopy. Significant increase in elastic modulus observed in 1 and 3mg ml^-1^ at 96 hrs. *indicates statistical significance (p<0.05).

### Changes in Cellular Morphometry due to Extracellular Density

Apparent changes in cell morphology were observed among all three collagen concentrations quantified across different planes of the microgel construct (**Fig. 3a**). Confocal imaging of ‘live microgels’ revealed differences in shape factor (SF), surface area to volume ratio (SAV) and longest axis of cells with microgels. SF defines the roundedness or stretched morphology of the cells, an SF of 0 indicates that a cell is completely stretched. A SF of 1 is assigned to a perfectly rounded cell. Cell shape factor index analysis showed a variable cell shape factor index in 1mg ml^-1^(0.45-0.6) and 3mg ml^-1^(0.3-0.55) microgels. A consistent cell shape was observed across all levels in 2mg ml^-1^ where the cells assumed a shape factor (0.6-0.62) which was significantly different from 1mg ml^-1^ and 3mg ml^-1^ (**Fig. 3b**). The cells embedded within 2mg ml^-1^ microgels also exhibited a consistent surface area to volume ratio (0.35-0.4) at all three planes across the microgel (**Fig. 3c**). Cell alignment on its longest axis in a plane is an apolar event that precedes cell division. The longest axes of the cells were significantly different between the microgel conditions. As cell alignment also correlates to cell shape, the cell alignment was shorter in the center and the bottom plane in 2mg ml^-1^ compared to 1mg ml^-1^ and 3mg ml^-1^. Overall, higher number of cells had the longest axis on the periphery of the microgel compared to that of the centre (**SI Appendix, SI Fig. S2c**).

**Figure 3.**
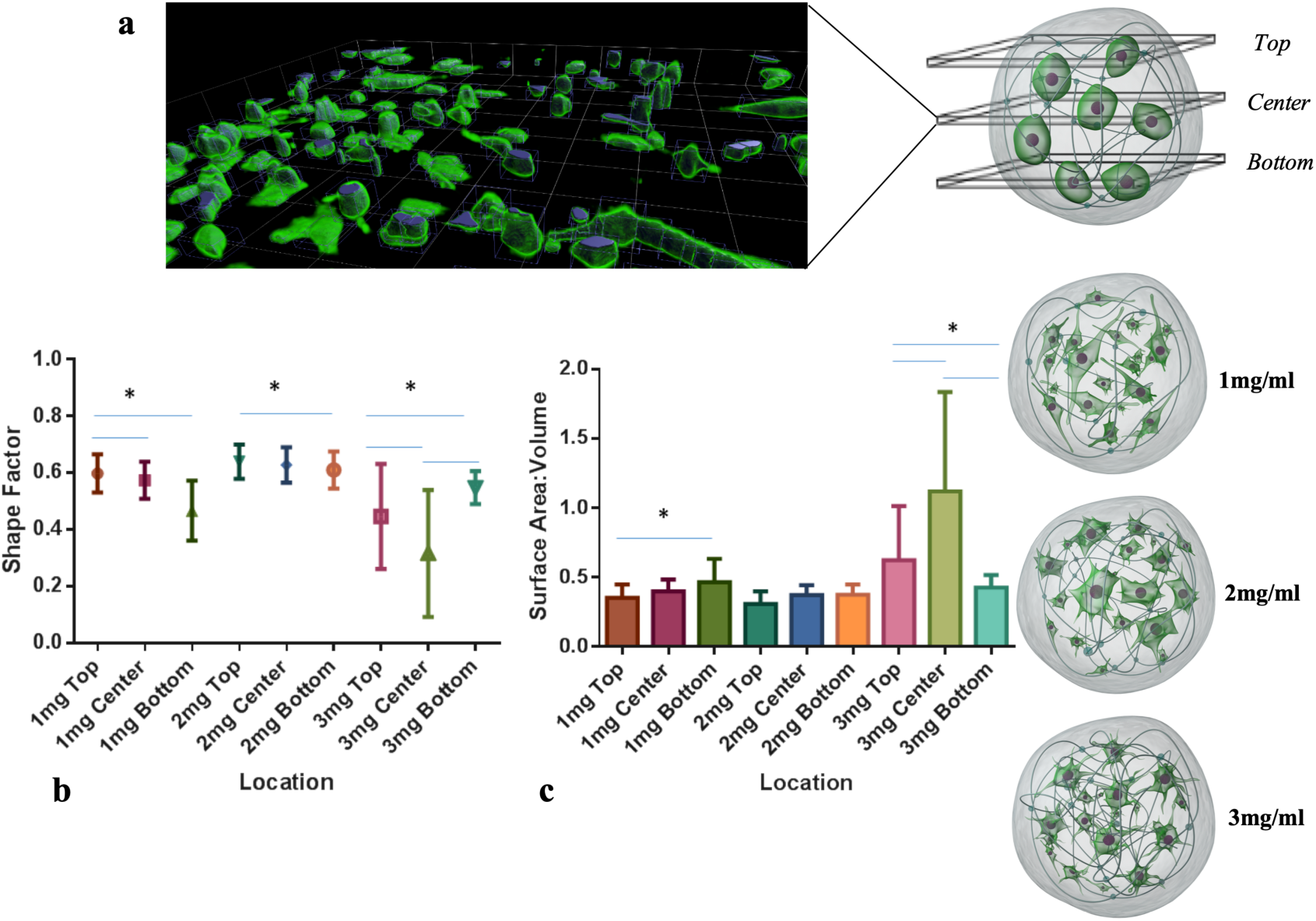
Microgel macromolecular concentration influences cell morphology, surface area to volume ratio and cell polarity. (a) 3D rendered image slice of a microgel acquired on a confocal microscope through 800µm depth with images taken 5µm apart, cells were stained by Calcien AM and visualised by an excitation wavelength of 488nm. (panels) representative images showing morphological differences at 96 hours in 1, 2 and 3mg ml^-1^ microgels (b) Shape factor (SF) of hMSCs measured across 3 planes (top, centre and bottom) on a microgel, shape factor ranging from 0 (rounded) to 1 (wide stretched) (c) Surface area to volume ratio (SAV) of cells across three levels. (c) Surface area to volume ratio (SAV) of hMSCs measured across 3 planes (top, centre and bottom) in microgels (d) Longest axis of hMSCs on microgels indicative of cellular anisotropy. *indicates statistical significance (p<0.05).

### Formation of Intracellular Inclusions in hMSCs

Cells embedded in different microgel concentrations were analysed for changes in intracellular organelles and composition. The electron micrograph from microgels demonstrated a higher number of cytoplasmic inclusions in 2mg ml^-1^ microgels than that in 1mg ml^-1^ and 3mg ml^-1^ microgels over 96 hours (**SI Appendix, SI Fig. S2a**). A detailed analysis revealed a significant difference in the percentage volume fraction per unit cell area embedded in different microgel concentrations. At 24 hours, the cytoplasmic inclusions were not significantly different between the groups, with the percent volume of 0.01-0.02% per cell area. At 96 hours the percent volume fraction of inclusions in 2mg ml^-1^ microgels increased to 0.15% compared to 0.03% in 1mg ml^-1^ and 3mg ml^-1^ microgels (**SI Appendix, SI Fig. S2b**). Additionally, numerous stacked rough endoplasmic reticula can be observed in 2mg ml^-1^ microgels in contrast to irregular distended rough endoplasmic reticulum in 1mg ml^-1^ and 3mg ml^-1^microgels.

### Extracellular Matrix Concentration Dependent Modulation of Integrin and Extracellular Matrix Gene Expression

#### Comparison to Tissue Culture Plastic

Integrins and extracellular matrix associated genes were quantified using real-time PCR arrays. At 24 hours, 52 genes and at 96 hours 65 genes were up or downregulated in the microgel groups relative to those in tissue culture plastic **(SI Appendix, Fig. S4a)** (**SI Appendix, Table S1, Table S2**). Extracellular matrix genes such as COL1A1, COL4A2, COL5A1, COL8A1, COL11A1, COL12A1, COL14A1, COL15A1, LAMA2, LAMA3, LAMB1, LAMC1, FN1, VTN, VCAN, ECM1 were downregulated (p<0.05) and LAMB 3 upregulated (p<0.05) in all microgel groups at 24 hours (**Fig. 4a**). Matricellular and adhesion associated genes PECAM1, SGCE, SPARC, SPG7, THBS2, THBS3, VCAM1, ICAM1 were also significantly downregulated (p<0.05) in all microgel groups (**Fig. 4b**). At 96 hours, COL1A1, COL4A2 COL5A1, COL7A1, COL8A1, COL16A1, VTN, FN1, LAMC1 were significantly downregulated in the 3mg ml^-1^ microgels (p<0.05), and COL11A1, COL12A1, COL14A1, COL15A1 were downregulated in all microgel groups (**Fig. 4c**). Strikingly at 96 hours, PECAM1/CD31 was upregulated in 1mg ml^-1^ microgel (fold 3.8, p=0.02) and 2mg ml^-1^ microgels (fold 5.6, p=0.02). CLEC3B (fold 2.6, p=0.005) and SPP1 (fold=2.97, p=0.04) were also significantly upregulated in 2mg ml^-1^microgels. Matricellular and adhesion associated genes SGCE, SPARC, THBS2, THBS3, VCAM1 and CLEC3B remained significantly downregulated (p<0.05) in 3mg ml^-1^ microgels at 96 hours (**Fig. 4d**).

**Figure 4.**
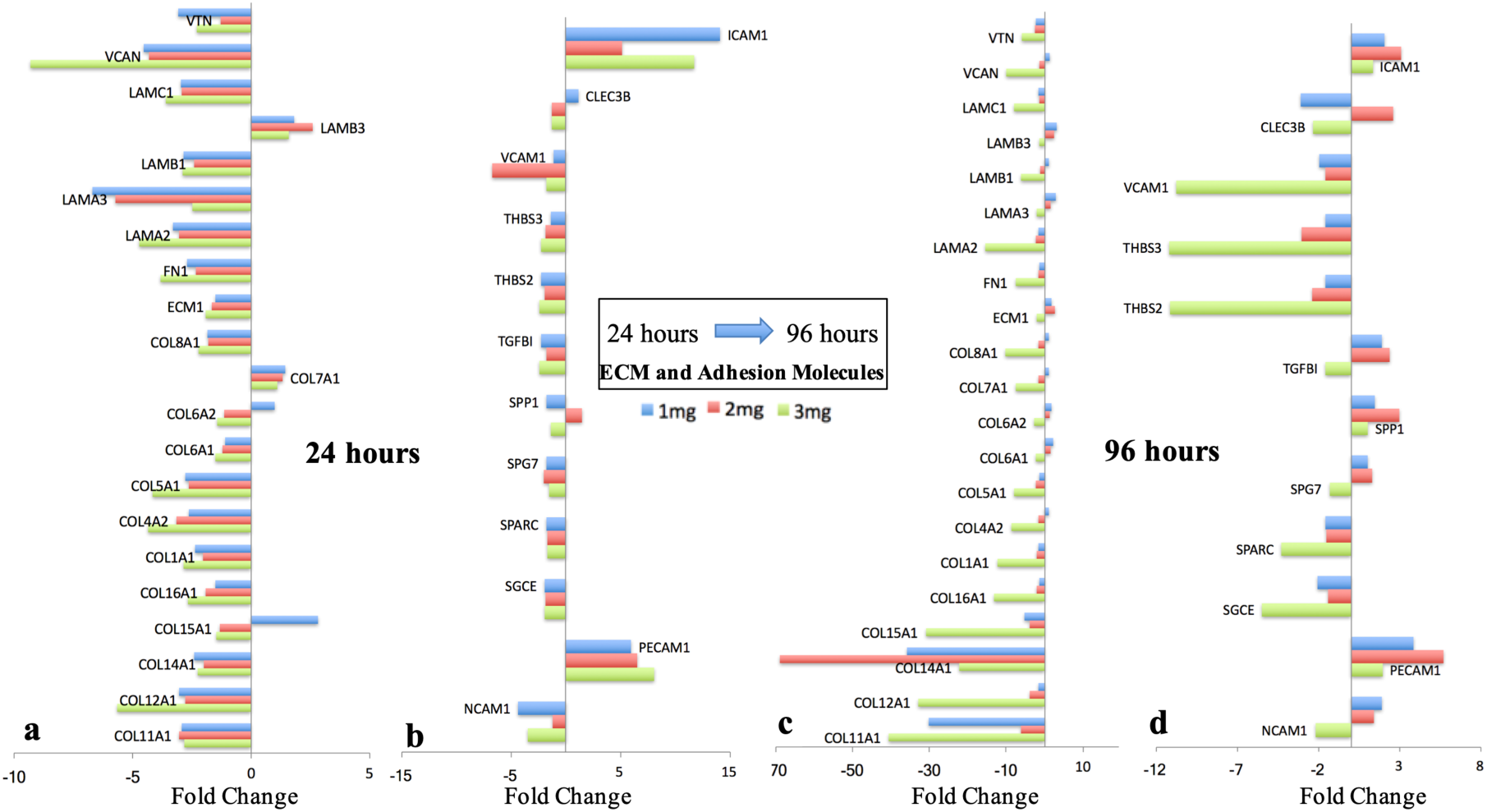
Increasing macromolecular concentration downregulates ECM and adhesion molecules gene expression. Early changes in (a) ECM genes (b) and adhesion molecules differentially expressed in microgel concentrations (1, 2 and 3mg ml^-1^) at 24 hrs relative to tissue culture plastic control. Late changes in (c) ECM genes (d) and adhesion molecules differentially expressed in microgel concentrations (1, 2 and 3mg ml^-1^) at 96 hrs relative to tissue culture plastic control. Fold change values are the mean level of differential expression (n=3/group) and based on a 1.5-fold threshold. 7

IPA analysis and canonical analysis showed similarities with respect to inhibition of several signaling pathways (**SI Appendix, SI Fig. S5a**) particularly integrin signaling (z score=<-2, p<0.05) at 24 hours in all microgel groups. Both in 1mg and 2mg ml^-1^ microgels activation of Rac, Rho GTPases, IL-8 was observed. A higher activation of the integrin signaling pathway in 2mg ml^-1^ microgels was seen due to a higher fold expression of integrins (**SI Appendix, SI Fig. S5a, b**) in the signaling cascade. Furthermore, functional analysis comparison (**SI Appendix, SI Fig. S5c**) exhibited several cell functions. These included inhibition of adhesion of extracellular matrix, cell spreading, differentiation and proliferation at 24 hours. Notable differences were observed at 96 hours between the 1mg, 2mg and 3mg ml^-1^ microgel groups with respect to functions such as proliferation, adhesion of extracellular matrix, angiogenesis and vasculogenesis. These functions were activated relatively higher in the 2mg ml^-1^ microgels, followed by 1 mg ml^-1^microgels and inhibited in 3mg ml^-1^ microgels. Correlation analysis on gene expression in 2mg ml^-1^ microgels showed a strong (Pearson’s P>0.6) positive correlation between 17 markers (**SI Appendix, SI Fig. S5d**). Integrins ITGA1, ITGA3, ITGA4, ITGA6, ITGA7, ITGB1, ITGB2 and ITGAV were significantly downregulated (p<0.05) in all microgel groups at 24 hours (**Fig. 5a**). Similarly, matrix metalloproteases such as MMP2, MMP16 and TIMP2 were downregulated, whereas MMP1 and MMP14 were significantly upregulated (p<0.05) (**Fig. 5a**). At 96 hours, integrin ITGA1 was observed to be significantly downregulated in all microgel groups (p<0.05). ITGA2 was upregulated in 1mg ml^-1^ microgels (fold 2.99, p=0.03) and 2mg ml^-1^ microgels (fold 3.57, p=0.026). ITGA5(fold 2.03, p=0.02), ITGB3 (fold 2, p=0.03) and ITGB5(fold 2.21, p=0.01) were significantly upregulated in 2mg ml^-1^ microgels (**Fig. 5b**). Strong Pearson’s correlation (P>0.6) between integrins ITGA2, ITGA4, ITGA5, ITGA6, ITGA7, ITGA8, ITGAV, ITGB3 and ITGB5 were also observed **(SI Appendix, SI Fig. S5e)**.

**Figure 5.**
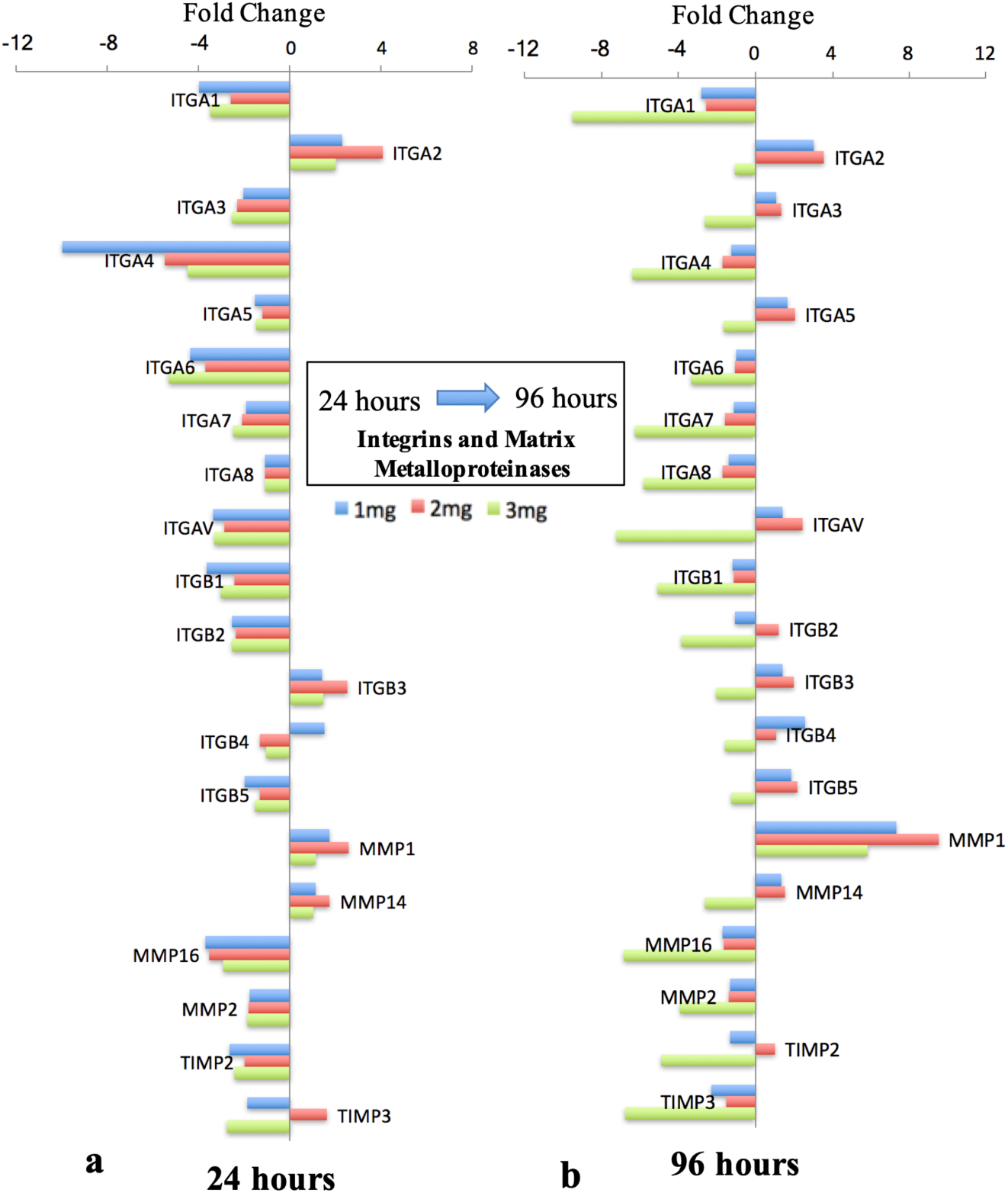
Microgel macromolecular concentration induces temporal changes in integrin gene expression. **(a)** 24 hrs **(b)** 96 hrs profile of cellular integrin molecules differentially expressed in microgel concentrations (1, 2 and 3mg ml^-1^) relative to tissue culture plastic control. Fold change values are the mean level of differential expression (n=3/group) and based on a 1.5-fold threshold.

#### Differences between Microgel Concentrations

At 24 hours, most ECM genes were downregulated across all microgel groups. However, gene expression levels of VTN, LAMA3, ITGA4, NCAM1 were significantly downregulated (fold <-1.5), and COL15A1, ICAM1, ITGB4 were upregulated (fold >1.5) in 1mg ml^-1^ microgel compared to 2mg ml^-1^ microgel and 3mg ml^-1^ microgel. Gene VCAM1 was significantly downregulated (fold <-1.5) and LAMB3, ITGA2, ITGB3 were upregulated (fold >1.5) in 2mg ml^-1^ microgels compared to 1mg ml^-1^ and 3mg ml^-1^ microgels. Genes COL4A2, COL12A1, VCAN, FN1, LAMA2 were significantly downregulated and PECAM1 upregulated in 3mg ml^-1^ microgels, compared to 1mg ml^-1^ and 2mg ml^-1^microgels. At 96 hours, upregulation (fold > 1.5) of genes COL6A1, COL6A2, LAMA3, LAMB3, NCAM1, and ITGB4 were observed to be significantly higher in 1mg ml^-1^ microgels than in 2 mg ml^-1^ and 3mg ml^-1^ microgels. Genes ITGA2, ITGA5, ITGAV, ITGB3, ITGB5, MMP1, SPP1, TGFB1, CLEC3B, ECM1, LAMB3, PECAM1, ICAM1 were observed to be upregulated (fold >1.5) in 2mg ml^-1^ compared to 1mg ml^-1^ and 3mg ml^-1^ microgels. Strikingly, 34 genes (fold <-1.5) [COL1A1, COL5A1, COL6A1, COL6A2, COL7A1, COL8A1, COL11A1, COL12A1, COL15A1, COL16A1, VTN, ECM1, FN1, LAMA2, LAMA3, LAMB1, LAMC1, ITGA1, ITGA3, ITGA4, ITGA5, ITGA6, ITGA7, ITGA8, ITGAV, ITGB1, ITGB3, NCAM1, SGCE, SPARC, TGFBI, THBS2, THBS3, VCAM1] were significantly downregulated in 3mg ml^-1^ microgels compared to those in 1mg ml^-1^ and 2mg ml^-1^ microgels.

### Improved Limb Salvage in a Severe Hindlimb Ischemia Model

The therapeutic efficacy of hMSC embedded microgels characterised based on previously quantified angiogenic paracrine effects (19) was tested in a recently developed hindlimb ischemia model (21). Microgels of 1mg ml^-1^, 2mg ml^-1^, 3mg ml^-1^ collagen concentration at a cell density of 8×10^5^ were cultured for 96 hours and were labelled with PkH26 dye prior to *in vivo* implantation. Approximately, 30±5 microgels were enumerated for implantation to optimally fill the perimuscular space, at the proximal site of ligation above the profunda femoris in the left limb. The right limb served as the contralateral non-occluded control. After the induction of double ligation (**Fig. 6a**), the animals received either (i) microgels alone (ii) 1mg ml^-1^ microgels with hMSCs (iii) 2mg ml^-1^ microgels with hMSCs (iv) 3mg ml^-1^ microgels with hMSCs (v) hMSCs at high-cell density (vi) hMSCs at low-cell density or (vii) saline (n=12/group). Laser Doppler imaging revealed improved blood perfusion in 2mg ml^-1^ microgels with cells as early as day seven with 60 ± 17% recovery at day 21 which was significantly higher than that compared to the control groups (Saline: 29± 5%, 1mg ml^-1^: 40.5± 10% 3mg ml^-1^: 43± 12%, 1 million cells: 37±10%, 50K cells: 37.5±8%, Microgels alone: 41±12%) (**Fig. 6b**). The ambulatory improvement and extent of foot necrosis were quantified using a modified Tarlov scale (22) and ischemia scale (23). A higher score indicates impaired use of the ischemic limb, severe tissue necrosis or autoamputation of toes. Only 10% of the animals did not develop any necrosis in the saline group compared to 16% in the microgels alone group, 30% in low-cell density group, 48% in high-cell density group, 25% in 1mg ml^-1^, 50% in 2 mg ml^-1^microgels with cells respectively (**SI Appendix, SI Fig. S6a**). However, only 15% of the animals developed severe necrosis or underwent autoamputation of toes; whereas severe necrosis was observed in 58% of animals treated with saline, 49% with low-cell density, 33% in high-cell density, 33% in 3mg ml^-1^ and 25% in 1mg ml^-1^ respectively. Microgels with cell groups showed no significant difference in ambulatory scores up to two weeks. But at the end of three weeks, a significant difference was observed between 2mg ml^-1^ microgels with cells compared to the saline, microgel alone and 3mg ml^-1^ groups (**SI Appendix, SI Fig. S6b**).

**Figure 6.**
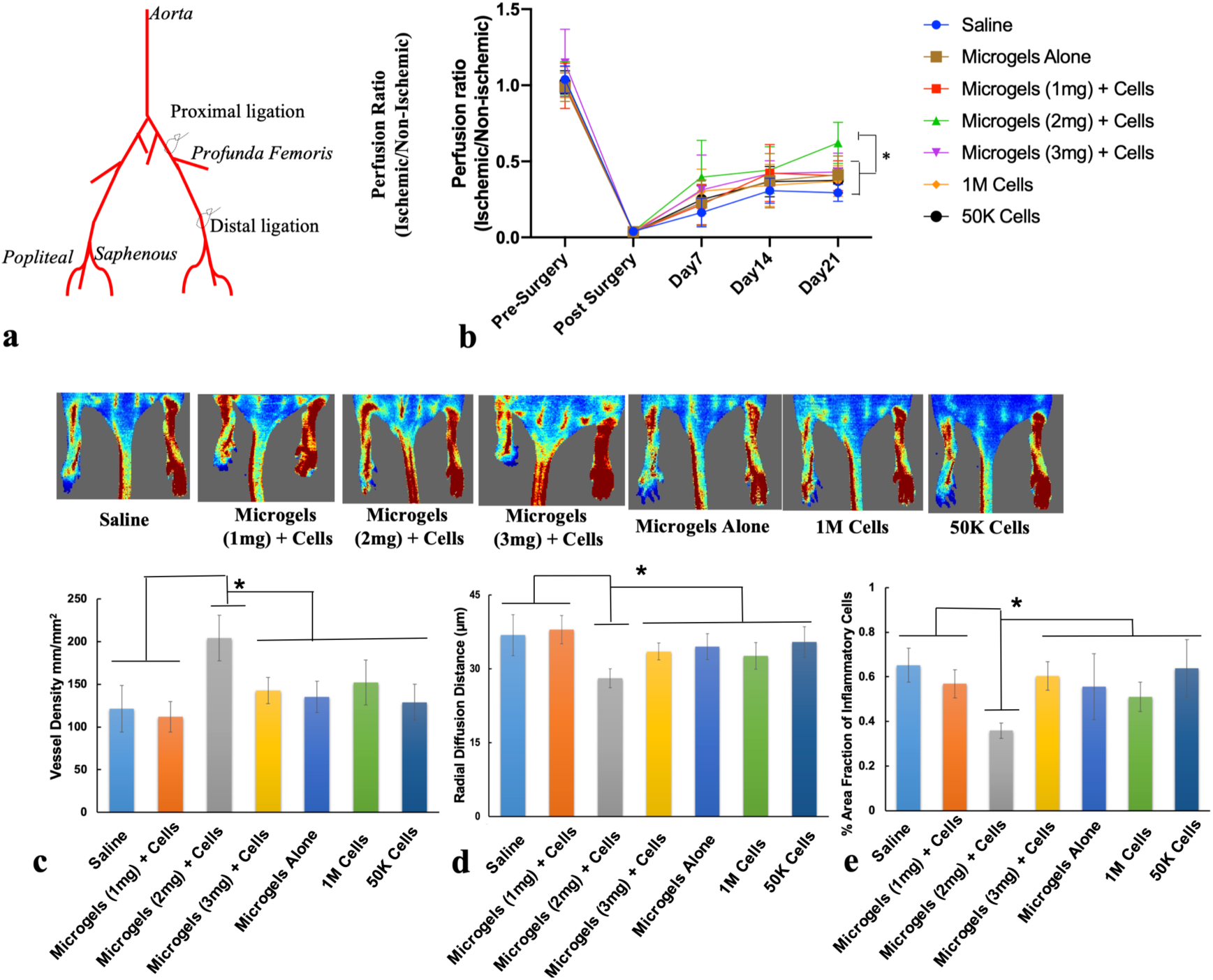
Improved blood flow in mice treated with hMSC embedded in microgels at a low-cell dose. (a) Schematic shows the double ligation sites on the femoral artery in the hindimb of Balb/c nude mouse model. (b) Laser Doppler evaluation of the ischemic (left) and non-ischemic (right) hindlimbs at day 21 (n=12/group, p<0.05). (c) hMSC embedded 2mg ml^-1^ microgels showed significantly higher capillary density and (d) a lower radial diffusion distance compared to control groups at day 21. (e) Significantly lower infiltration of inflammatory cells was observed in 2mg ml^-1^ microgels with hMSCs compared to rest of the treatment groups. (n=12/group, p<0.05), Scale bar 50μm. *indicates statistical significance (p<0.05).

### Microgels Embedded with hMSCs Enhanced Angiogenesis and Reduced Inflammatory Response

To confirm the therapeutic angiogenesis effect of microgels embedded with low-dose cells, the muscle tissues from the gastrocnemius and lower quadriceps were subjected to histological analyses. At day 21 2mg ml^-1^microgels with hMSCs were found to have higher capillary density (**Fig. 6c**) (204±26 capillaries/mm^2^) compared to microgels alone (135±18 capillaries/mm^2^), 1mg ml^-1^ microgels (111±17 capillaries/mm^2^), 3mg ml^-1^ (142±15 capillaries/mm^2^), high-cell density (152±26 capillaries/mm^2^), low-cell density (129±21 capillaries/mm^2^) and saline (121±27 capillaries/mm^2^) groups respectively. A significantly lower radial diffusion (**Fig. 6d**) distance was observed in 2mg ml^-1^microgels with cells (28μm) compared to that of controls, which is a measure of the distance between the capillaries. The shorter the distance, the higher is the degree of tissue perfusion. Subsequent antibody staining for the endothelial marker CD31 (PECAM-1) (**SI Appendix, SI Fig. S7**) confirmed the presence of a higher number of blood vessels in the hMSC embedded microgels than that seen in the controls. In addition, a significant reduction in the infiltration of inflammatory cells was observed (**Fig. 6e**) in the microgels with hMSCs group (0.35±0.03) compared to microgels alone (0.55±0.13), 1mg ml^-1^ microgels (0.56±0.06), 3mg ml^-1^(0.60±0.06) high-cell density (0.50±0.06), low-cell density (0.63 ±0.12) and saline (0.65±0.07). A significant difference was also observed in the high-cell density group compared to the low-cell density and saline groups. Immunostaining for CD68 (**SI Appendix, SI Fig.S7**), a macrophage marker, further confirmed the reduced inflammation in 2mg ml^-1^ microgels with hMSCs compared to the control cell and microgel groups.

### Microgels Embedded with hMSCs Enhanced Angiogenesis Factor Expression in Muscle Tissues

Quantitative analysis of cytokines and chemokines from muscle tissue extracts showed significantly higher levels of proangiogenic factors (ANG-2, sCD31, Endoglin, FGF-2, Prolactin, VEGF-C) in the muscle treated with microgel embedded hMSCs than those of saline (**Fig. 7a, b**). Significant differences were also observed in ANG-2, sCD31 and Prolactin in the muscle tissue between 2mg ml^-1^microgel embedded hMSCs and high-cell density treatment. Factors ANG-2, sCD31, Endoglin, FGF-2, Prolactin and VEGF-C were found to have lower levels of expression in the low-cell density and microgels alone groups than those of 2mg ml^-1^microgel embedded hMSCs. Likewise, the gene expression level of ANG-1 (fold 4.31), FGF2 (fold 5.05), HIF-1α (fold 2.23), IL-6 (fold 3), PGF (fold 6.04), VEGF-A (fold 3.76), PDGFR-A (fold 2.89), ICAM-1 (fold 4.07), PECAM-1 (fold 6.71), MMP-9 (fold 3.85) was found to be significantly higher (p<0.05) (**SI Appendix, SI Fig. 8a, b**) in the muscle tissues treated with hMSC embedded 2mg ml^-1^ microgels than that in the control cell and microgel groups. Canonical pathway analysis showed activation of several pathways such as eNOS, IL-6, IL-8, ILK, integrin, NF-kB, nitric oxide, PI3K/AKT and VEGF signaling (**Fig. 8a**). A higher activation (z score >2, p<0.05) in IL-8, PI3K/AKT and VEGF signaling was seen in tissues from hMSC embedded 2mg ml^-1^microgels compared with the high-cell density group. Both low-cell density and microgels alone treatment led to inhibition of the canonical pathways. Further, functional analysis comparison (**Fig. 8b**) revealed activation of several cell functions associated with angiogenesis, adhesion of vascular endothelial cells, growth of blood vessel, and inhibition of cellular apoptosis and reduced infiltration of macrophages and antigen presenting cells in the hMSC embedded microgel group.

**Figure 7.**
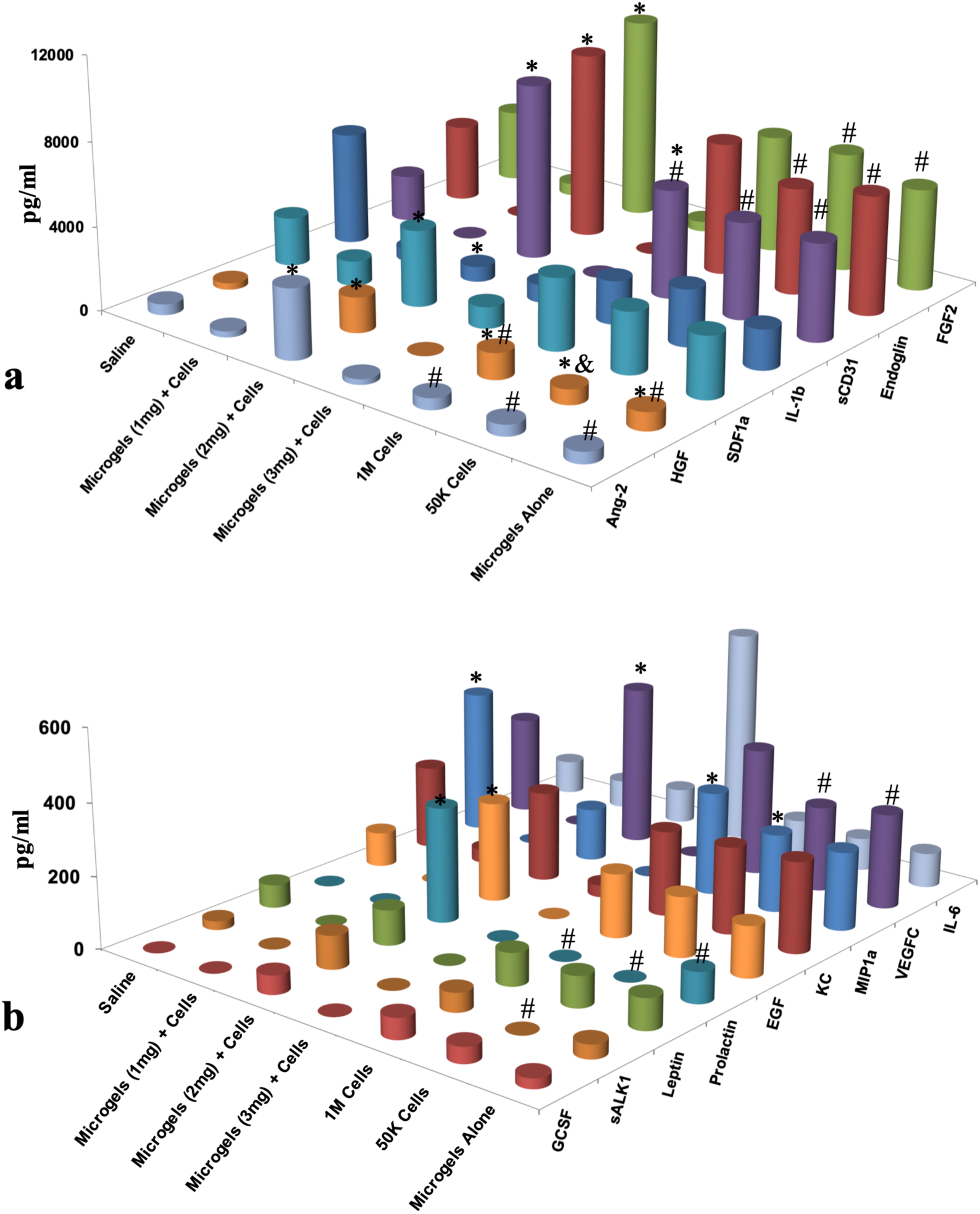
High levels of pro-angiogenic markers (sCD31, ANG-2 FGF-2, Endoglin, EGF, HGF, SDF-1a, Prolactin, VEGF) (a) and (b) were measured in 2mg compared 1, 3mg groups and control groups in the tissues at day 21. Statistical significance tested with two-way ANOVA, p<0.05; (n=3/group; n=1 pooled from 4 animals). *#& indicates statistical significance (p<0.05).

**Figure 8.**
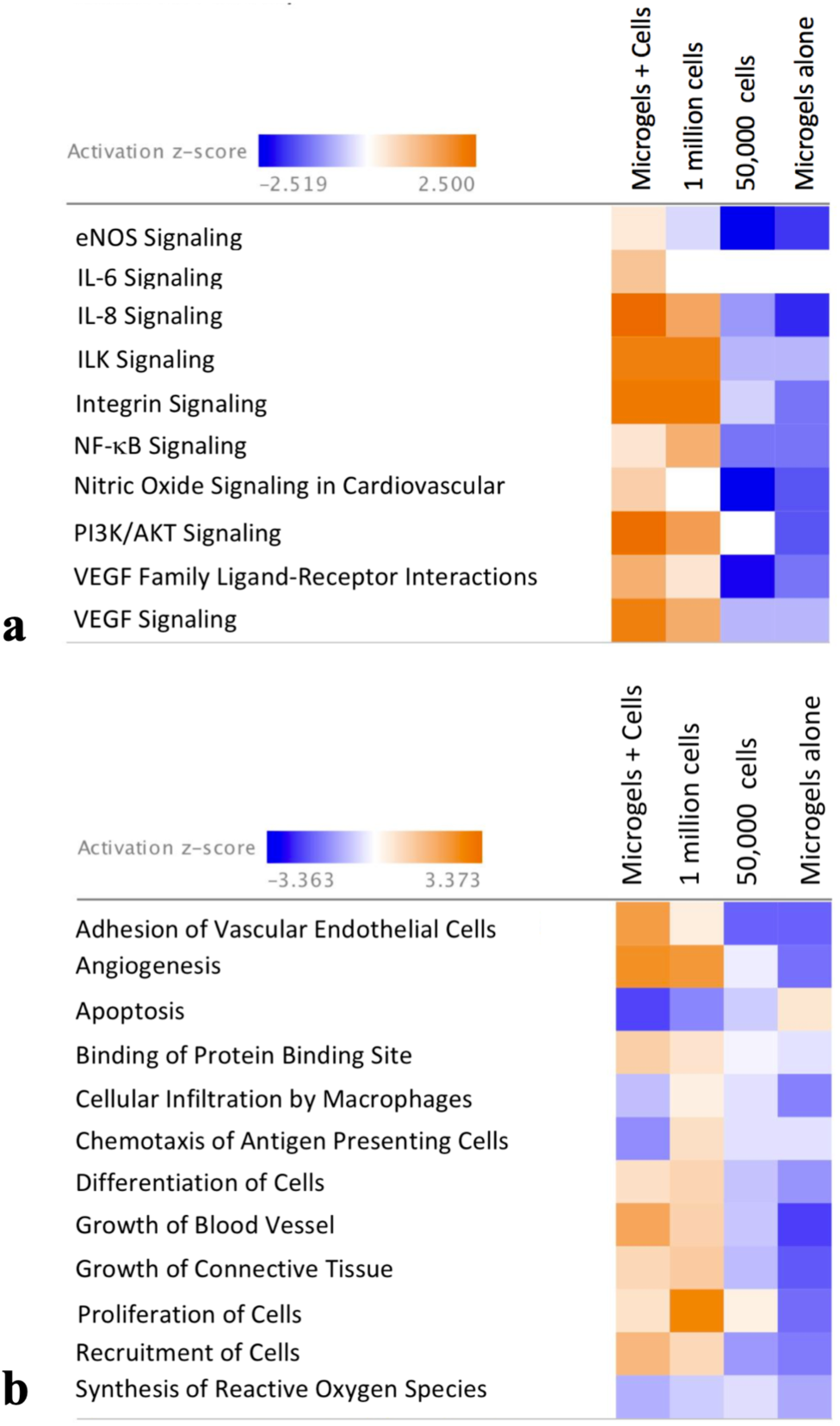
Pathways affected in mice treated with human mesenchymal stem cells embedded in microgels after hindlimb ischemia. (a) Heatmap of z-scores (indicating activation or inhibition) of canonical pathways Ingenuity Pathway Analysis (IPA) with a threshold of 1.3-fold expression, p<0.05. The dataset represents changes in gene expression in treatment groups: microgels with hMSCs (2mg ml^-1^), 1 million hMSCs, 50, 000 hMSCs and microgels alone (b) Heat map displaying z-scores of biological functions (IPA) between treatment groups at day 21.

### Differences in Tissue *N*-Glycan Distribution Treated with Microgels Embedded with hMSCs

To obtain the *N*-glycan population in tissues, unfixed muscle cryosections were treated with dialysed glycerol-free PNGase F and the deposition of ionising matrix 2,5-DHB, followed by MALDI-TOF MS analysis in imaging mode. **Figure 9a** shows a profile mass spectrum of muscle sections from the upper gastrocnemius in the treatment groups with normalised intensities for *m/z* 1101± 2 and *m/z* 2135± 3, and consecutive H&E stained muscle sections. The *N*-glycan candidates were chosen for their high abundance and differential distribution in the group treated with hMSC embedded 2mg ml^-1^ microgels. To confirm the identity of the *N*-glycans, supernatants from microdissected and enzyme treated tissues, the samples were analysed on a MALDI-TOF/TOF instrument. Tissue supernatants obtained from in-solution digests (**Fig. 9b**) also contained high *S/N* peaks corresponding to *m/z* 1101 and *m/z* 2135. GlycoMod prediction indicates Hex_4_(Deoxyhexose)_1_(Pent)_2_ to be the only possible glycoform (glycoform mass 1058.3, Δmass 1.3 Da). Hex_4_(HexNAc)_2_(Deoxyhexose)_1_ + (Man)_3_(GlcNAc)_2_ (glycoform mass 2092.8, Δmass 1.3 Da) or Hex_1_(HexNAc)_1_(Deoxyhexose)_3_(Pent)_3_ + (Man)_3_(GlcNAc)_2_ (glycoform mass 2091.75, Δmass 2.25 Da) are the most likely glycol forms predicted for *m/z* 2135±3. Both structures correlate with the chitobiose core structure of N-glycans. Other structures were ruled out according to mass accuracies and fragment ions generated in MS/MS experiments confirming the ions to be carbohydrates.

**Figure 9.**
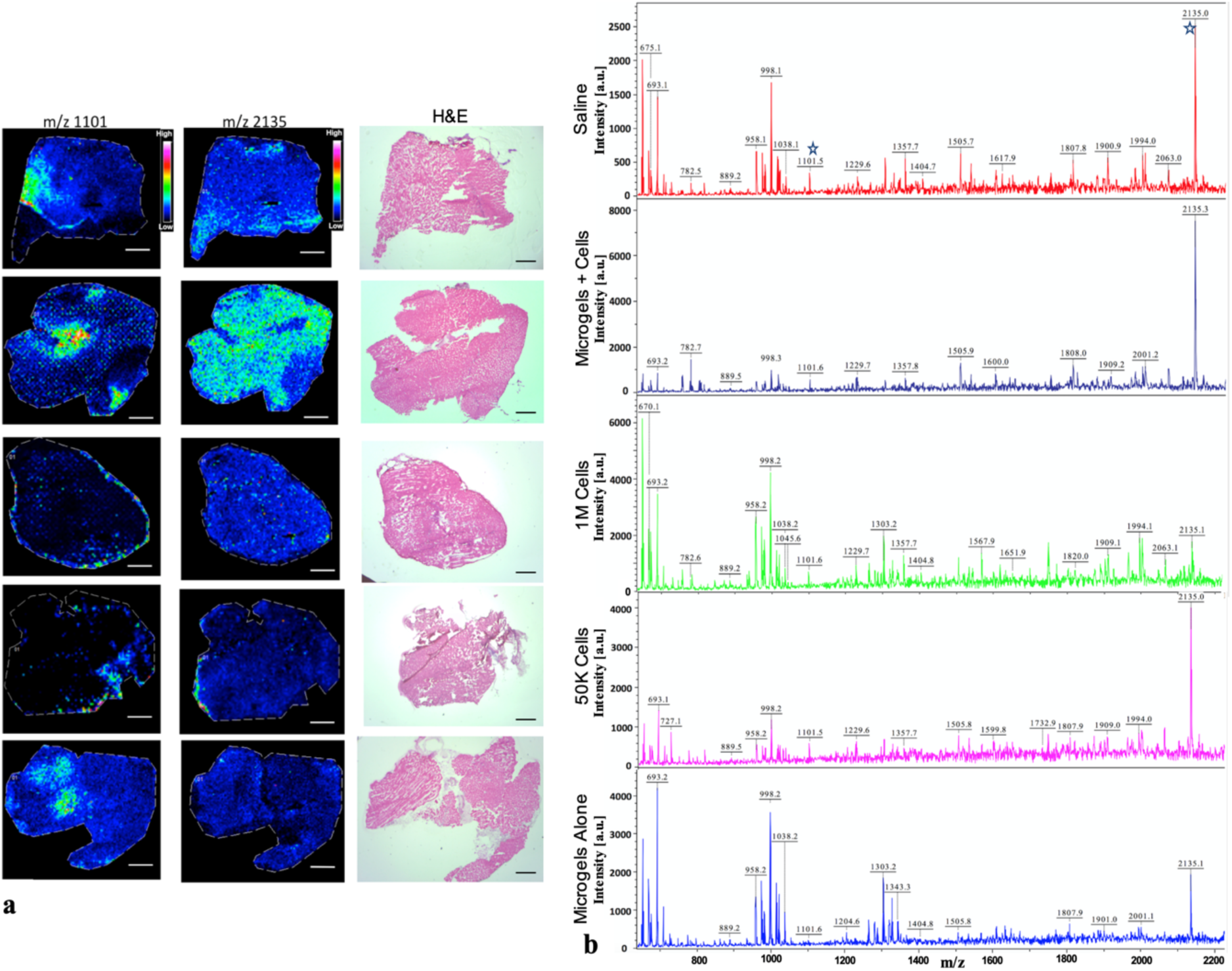
In situ profiling of N-linked glycans from tissues. (a) Ion intensity maps of N-glycans showing distribution of m/z 1101 and m/z 2135 and consecutive H&E stained muscle sections. (b) In-solution tissue digest of tissue samples showing ions identified from the MADLI/IMS experiments. Ion intensities are normalised to the total ion current for each ion across the tissue and rainbow scale bars indicate the range of intensities plotted.

### Higher Expression of *N*-Linked Sugars in Microgels with hMSCs Treatment Group

To validate the presence of predominant *N*-linked sugars predicted by mass spectrometry, lectin histochemistry assay was performed. Expression of N-acetyl-D-glucosamine and sialic acid was confirmed with WGA lectin. Presence of fucosylation was confirmed by lectin UEA-I that has a binding specificity to α-(1 → 2)- and α-(1 → 6)-linked fucose. WGA binding was observed to be significantly higher in 2mg ml^-1^microgels with hMSCs treatment group compared to the controls, except 1 million cell group (**SI Appendix, SI Fig. S10 a, b**). Whereas UEA-I expression was significantly higher (**SI Appendix, SI Fig. S10 a, c**) in 2mg ml^-1^ microgels with hMSCs compared to control groups.

## Discussion

The effect of cell-material interactions in a physiologically similar 3-D environment has a tremendous impact on cell behaviour. Engineering a tissue-specific extracellular environment through modulation of macromolecular concentration replicates the different stages of tissue repair and remodelling. The use of naturally occurring extracellular matrices innately provides instructive cues which helps maximise cell-cell and cell-matrix interactions. Use of tailored matrices as a cell delivery platform not only allows high cell retention and survival but also serves as a pre-conditioning or priming platform. In the present study, the microgel platform previously characterised (19), was tested for inducing therapeutic angiogenesis in a severe hindlimb ischemia model. It has been demonstrated that hMSCs delivered via gelatin microcryogels at a cell density of 1 ×10^5^ cell dose have a better angiogenic effect than 1 ×10^5^ cells delivered on their own (24). Additionally, a dose-dependent study confirmed the efficacy of 1 ×10^5^ hMSCS from both CLI and normal donors and showed equivalent neovascularisation in a murine hindlimb ischemia model (25). A further two-fold lower cell dose (5×10^4^ hMSCs) primed with a growth factor cocktail containing bFGF, PDGF-AA and heregulin-β1 for eight days was shown to be effective in inducing angiogenesis *in vivo* (26). Several cell densities (0.1 million, 0.4 million and 0.8 million cells ml^-1^) were tested previously(19), and the cell density used in the current study was determined based on the high-level of paracrine output observed on the microgel platform. As a first, in our study, we demonstrate the *in vivo* efficacy of hMSCs at a dose of 5 ×10^4^ embedded in type-I collagen microgels primed *in vitro* for 96 hours without addition of any external growth factors.

In a physiological state, the 3-D extracellular environment influences cell behaviour including proliferation, matrix metalloproteinase mediated migration, expression of cell-adhesion integrins, all of which are important events for angiogenesis. Cells grown in a 3-D microgel environment retained phenotypic cell surface markers (**SI Appendix, SI Fig. S1e)** with high cell survival with over 80% viability observed over two weeks (**SI Appendix, SI Fig. S1f**). Genotypic analyses have revealed 3-D cell aggregates (27) or cells grown in 3-D hydrogels (28) in the presence of ECM proteins are more relevant to *in vivo*, as compared to cells grown on 2-D. 3-D microgels developed here promote both cell-cell and cell-matrix interactions (**SI Appendix, SI Fig. S1b, c, d**). In our previous work, a reduction in the proliferative rate on microgels was shown to be approximately four-fold than that of tissue culture plastic with a higher expression of matrix metalloproteinases(19). Several others in the field have demonstrated that crosslinking the collagen matrix enhances angiogenesis, whereas a higher matrix density has a negative effect on angiogenesis (29, 30). Notably, both the current and previous work employs optimal crosslinking with varying collagen concentrations. The results show that changes in matrix stiffness over-time is collagen concentration dependent and this is reflected in sensing stiffness at the cellular level. The subcellular localisation of transcription factor YAP/TAZ is tightly regulated by cell substrate rigidity that results in actin cytoskeleton remodelling (31). Hence YAP/TAZ serves as molecular mechanosensors, where its translocation to the nucleus affects cell migration and proliferation (32).

In a 3-D environment, cells secrete ECM to provide structural and biochemical support, and the remodelling of ECM is dependent on the activity of YAP (33). These results indicate that significant ECM remodelling in 1mg ml^-1^ and 3mg ml^-1^ microgels corresponds to higher nuclear translocation of YAP/TAZ and an increase in matrix stiffness greater than 1kPa. Although, an increase in modulus is observed in 2mg ml^-1^ microgels at 96 hours, this change is significantly lower than for 1mg ml^-1^ and 3mg ml^-1^ microgels. A recent study concluded that YAP/TAZ activity was not regulated by stiffness alone, but other factors such as cell spreading permissive environment and generation of contractility that leads to activation of YAP/TAZ complex(34). Our data clearly demonstrated that the increase in macromolecular concentration restricts cell spreading, and the distinct YAP/TAZ nuclear localisation in 1 and 3mg ml^-1^ microgels compared to 2mg ml^-1^ could be due to the dynamic changes in MSC contractility and onset of stress induction as previously reported (34, 35). The microgel constructs were also observed to contract in size due to MMP-1 activity by the MSCs. The rate of contraction was dependent on cell density and collagen concentration, with overall up to 30% reduction in construct size (19). To further explore the influence of biophysical factors that decouple cellular contraction, we investigated YAP/TAZ translocation after inhibition of contractility with blebbistatin during the pre-conditioning period. Nuclear YAP/TAZ co-localisation was not significantly affected in 1 and 3mg ml^-1^ compared to controls (**SI Appendix, SI Fig. S3c)**. Corroborated with changes in cellular spreading there was a significant increase in nuclear YAP/TAZ in 2mg ml^-1^ microgels, suggesting limited matrix degradation and inter-cellular tractions due to cell-cell contact may play an important role in local deformative mechanical changes(36) that is independent of measured bulk hydrogel stiffness.

Further, previous studies have looked at the potential of mesenchymal stem cells in forming interconnected endothelial cell-like networks, exploring a potential trans-differentiation mechanism. However, no studies to date have looked at morphological changes and paracrine effect of MSCs in a three-dimensional environment with maintenance of multipotency. Cell-mediated ECM changes in 1mg ml^-1^ and 3mg ml^-1^ microgels had a significant impact on the cell morphology throughout the microgel construct; whereas a more consistent cell morphology is observed in 2mg ml^-1^ microgels. Additionally, the surface area to volume ratio of cells in 1mg ml^-1^ and 3mg ml^-1^ microgels showed differences in distributions of bigger and smaller cells through the microgel. A consistent and smaller surface to area volume ratio observed in 2mg ml^-1^ microgels is indicative of a high metabolic activity due to the shorter transport distance of the metabolites (37).

Local confinement within a 3-D matrix and cell shape dictate cell alignment on its longest axis prior to division is determined by (38). In the current study, the longest axis of the cells was measured to be under 80μm, which is lower than that of the MSCs on polystyrene surfaces where cells spread freely, and elongation of the cell along its longest axis and higher anisotropy dictates the rate of cell division (39). The presence of abundant dense inclusion bodies within cells embedded in 2mg ml^-1^ microgels, distinct from the intracellular organelles, suggests a secretory cell behaviour (40-42). Although the exact nature of the inclusions were not characterised in the study, the secretory products may correlated to the paracrine response previously reported (19). Pearson’s correlation analysis showed strong positive correlations of molecular markers to CLEC3B in the integrin profiling experiment. CLEC3B (C-type lectin domain family 3, member b) or Tetranectin is known to encode a protein involved in packaging molecules destined for exocytosis (43, 44).

In the current study, the changes in the ECM composition during the pre-conditioning period was not investigated. This is because the aim was not to replicate the complexity of the hMSCs bone marrow ECM ‘niche’, but the pre-conditioning effects of microgel concentrations that synergistically modulate integrin expression and paracrine signaling toward a pro-angiogenic phenotype in hMSCs. Over 80% of ECM and cell adhesion associated genes were altered at 96 hours in the microgels groups relative to tissue culture plastic. Integrin expression is modulated by “outside-in” signaling due to clustering and formation of focal adhesions linked to the cytoskeleton (45). The integrins ITGA1, ITGA2, ITGA4, ITGA5, ITGAV, ITGB1 and ITGB3 are strongly induced by proangiogenic mediators and growth factors (46, 47). A significant increase in ITGA2, ITGA5, ITGB3 and ITGB5 gene expression in the 2mg ml^-1^ microgel at 96 hours may be linked stronger integrin-cytoskeleton interactions within the local microenvironment. Particularly, integrin AVB3 and VEGFR2 is known to synergistically contribute to angiogenesis by forming a physical complex (48). At the protein level, modulation of integrin AVB3 and its ligand vitronectin, as a function of extracellular collagen concentration is further demonstrated on the microgel platform (**SI Appendix, SI Fig. S4b**,**c**). A strong positive correlation was also observed in the gene expression of integrins ITGA2, ITGA5, ITGA6, ITGB3 and ITGB5, among others. Additionally, the higher gene expression of ICAM-1 and PECAM-1 cell adhesion molecules is linked to the regulation of angiogenesis (**SI Appendix, SI Fig. S9e**) via several known mechanisms (49). However, additional perturbation studies need to be conducted to establish a direct association of integrins and angiogenic growth factor secretion in the microgel platform. Together, these findings suggest that synergetic expression of integrins and growth factor expression previously reported maybe linked to the proangiogenic response in mesenchymal cell types embedded in 3-D microgels of defined matrix density(30).

Finally, the *in vivo* therapeutic efficacy of the 2mg ml^-1^ microgels were demonstrated in a severe double-ligation model which manifests acute clinical signs of necrosis due to multiple occlusions in the femoral artery. Microgels with hMSCs resulted in limb salvage in 50% of the animals, higher than those of the saline, cell controls and microgel groups. Furthermore, significantly higher reperfusion was observed in animals treated with microgels with cells compared to that of the controls. Reperfusion resulting from neovascularisation was confirmed by histological evaluation of capillary density in the muscle tissues. Furthermore, muscles tissues were profiled for proangiogenic markers to investigate the presence of growth factors and cytokines involved in angiogenesis. The muscle tissue from mice that received primed hMSCs embedded in microgels showed higher levels of proangiogenic cytokines ANG-2, sCD31, Endoglin, FGF-2, Prolactin and VEGF-C, indicating that the enduring effect of the treatment contributes to functional angiogenesis via growth factor expression. Importantly, these observations were also demonstrated at the gene expression level. IPA analysis revealed 40 out of 68 genes had an expression direction consistent with increased angiogenesis in mice treated with hMSC embedded microgels (**SI Appendix, SI Fig. S9a-d**). Given the expression of growth factors as a ‘signature’ of regenerative tissue, it was hypothesised post-translational modifications, particularly differences in *N*-linked glycan expression, are induced by this pro-angiogenic stimulus. It has been reported that the endothelial cell glycome is sensitive to changes in cytokines and growth factors in the tissue (50). Some of these changes such as the extension of *N*-glycan structures are responsible for activating pro-angiogenic signaling pathways (50, 51). Although the protective role of *O*-linked glucosamine is known in neonatal cardiomyocytes after ischemia-reperfusion (52), little is known about the changes in *N*-linked glycans after reperfusion. Our data suggests, unique distribution and abundance of *N*-linked sugars in a pro-angiogenic environment induced by microgels with hMSCs. Furthermore, lectin histochemistry confirmed higher expression of sugar moieties which have been previously demonstrate to be essential in endothelial function and vascular stability(53-55). These results suggest that the microgels with cells treatment group may be responsible in the restoration endothelial glycocalyx and thereby improving the therapeutic effect. However, future studies are needed to delineate the mechanism(s) that are involved in the dynamic changes that occur in the glycome after induction of angiogenesis. Taken together, the study highlights the importance of hMSC priming, demonstrating a therapeutic effect with significantly lower cell-dose on the microgel platform.

## Conclusion

The results of our study demonstrate that MSCs embedded in a three-dimensional collagen microgel cell delivery platform induce temporal pro-angiogenic response by both cell permissive remodeling of the extracellular environment and matrix guided changes in cellular function via mechanotransduction. Systematic investigation of the biochemical nature of the ECMs during the preconditioning in different collagen concentrations and its mechanistic influence on over or under expression of cellular receptors will be the focus of our future study. From a cell therapy point-of-view, our model platform offers a significant benefit over other cell delivery platforms with the use of a twenty-fold lower cell dose than that of the gold-standard used in pre-clinical hindlimb ischemia studies. This attributes to the importance of preconditioning hMSCs on a 3-D microgel platform that enables administration of low-cell dose as a localised therapy to reverse ischemia. Furthermore, these findings will be appreciated as increasingly significant, as future studies will investigate ECM-based three-dimensional niches using our platform technology for engineering constructs that will allow replication of native cellular microenvironments for enhancing the regenerative capacity of stem cells.

## Materials and Methods

### Fabrication of Microgels

Microgels were fabricated as described previously (19, 20). Briefly, bovine tendon derived type-I atelocollagen (1/2/3mg ml^-1^ final) was neutralised in sodium hydroxide 1M and phosphate buffer saline 10X. To crosslink the microgels four-star poly(ethylene glycol) succinimidyl glutarate, Mw = 10,000 (4S-StarPEG) JenKem Technology U.S.A. (Allen, TX) was added in 1:1 molar ratios. Finally, hMSCs were added to the gel forming mixture at 0.8×10^6^ cells ml^-1^. The forming gel solution was then micro-dispensed as 2 μL droplets on a hydrophobic surface (commercial Teflon^®^ tape) to create a spherical shape and incubated for 40 minutes at 37°C for stable cross-linking (**SI Appendix, SI Fig. S1a**). Microgels were then transferred to 24-well plates containing hMSC media and incubated at 37°C and 5% CO2. For inhibition studies, a medium with inhibitor of blebbistatin at 10 μM concentration was added to different microgel concentrations. Following 96 hrs of pre-conditioning, the microgels were fixed and permeabilised for immunostaining.

### YAP/TAZ Localisation Analysis in hMSC Embedded in Microgels

Microgels were fixed in 4% paraformaldehyde for 30 minutes and permeabilised with 0.1% Triton X for two hours. Microgels were then incubated with Yes-associated protein (YAP) and TAZ (transcriptional coactivator with PDZ-binding motif) primary antibody (1:250) in 2% goat serum overnight at 4C. Alexafluor^®^ 488, Goat anti-mouse IgG (1:100) secondary antibody was incubated for one hour at room temperature in blocking buffer. Following PBS washes, nuclei were counterstained with Hoechst (Life Technologies) for five minutes. Samples were then imaged as Z-Stacks on Andor™ Olympus Spinning Disk Microscope (Andor, Belfast, Northern Ireland), using Andor™ IQ software, with an X 20 or X 40 oil immersion objective lens. Pearson’s co-localisation coefficients were calculated using Volocity^®^ 5.0 software (Perkin Elmer Inc., Waltham) (56).

### Microgel mRNA Extraction and ECM and Adhesion Molecule PCR Arrays

For *in vitro* experiments, total RNA from the microgels at 24 and 96 hours timepoints were preserved in TRI^®^ reagent (Sigma-Aldrich) followed by extraction using a RNAeasy^®^ Microkit (Qiagen GmbH, Germany). All RNA samples were treated with DNase to remove contaminating genomic DNA. RNA quality was assessed using the Agilent^®^ 2100 Bioanalyzer™ (Agilent Technologies, CA) and samples with RNA Intergrity Number (RIN) >7 were used for downstream cDNA conversion with RT^2^ first-strand kit (Qiagen GmbH, Germany) according to the manufacturer’s protocol. Eighty-four gene targets were analysed on human ECM and adhesion molecules RT^2^ profiler PCR array (PAHS-013Z, SaBiosciences Corp., USA) for 1, 2 and 3mg ml^-1^ cell embedded microgels at 24 and 96 hours n=3 per sample; with n=1 pooled 20-25 microgels from a single experiment.

### Multiplex Cytokine Analysis

Microdissected hindlimb muscle tissues were pooled into groups of three biological replicates (n=3) per treatment group and stored at −80°C in lysis buffer (Triton 1%, NaCl 0.1M, Tris HCl, pH 7.4) containing a cocktail of protease inhibitors. Tissues were homogenised using TissueLyser LT (Qiagen) for 30 minutes at 50Hz with intermittent cooling every ten minutes. Samples were centrifuged at 13,000 RPM for fifteen minutes. Total protein in the supernatant was quantified using a BCA protein assay (Pierce and Warriner) and the total concentration across all the samples was normalised. Tissue lysates were analysed for 24 analytes (Angiopoietin-2, G-CSF, sFasL, sAlk-1, Amphiregulin, Leptin, IL-1b, Betacellulin, EGF, IL-6, Endoglin, Endothelin-1, FGF-2, Follistatin, HGF, PECAM-1, IL-17, PLGF-2, KC, MCP-1, Prolactin, MIP-1α, SDF-1, VEGF-C, VEGF-D, VEGF-A, and TNFα.) using a MILLIPLEX^®^ MAP mouse angiogenesis magnetic bead-based assay (EMD Millipore). The assays were read on a calibrated Bio-Plex^®^ 200 System. Using Bio-Plex^®^ manager, the optimised standard curves were used to calculate the concentration of analytes. Measurements that showed a coefficient of variability of <20% were included in the analysis.

### Statistical Analyses

Statistical analyses were performed using GraphPad Prism^®^ Version 5 (USA) and IBM SPSS^®^ Version 21, data were compared using one-way or two-way analysis of variance (ANOVA) based on the number of factors analysed, followed by a Tukey post-hoc comparison test. Bivariate analysis was performed to identify strong (Pearson’s p>0.6) correlations. Statistical significance was set at p < 0.05 (*).

### Data Availability

All relevant data, associated protocols, and materials are within the paper and *SI Appendix.* If any additional information is needed, it will be available upon request from the corresponding author.

## Acknowledgements

This work is supported by Science Foundation Ireland (SFI) and is co-funded by the ERDF under Grant Number [13/RC/2073] and Grant No. [09/SRC/B1794]. Research at M.M-D’s laboratory was supported by EMBO short-term fellowships [ASTF-236-2015] and COST action [BM1104]. We thank Mr. Matthias Holzlechner and Dr. Ernst Pittenauer for their valuable support and discussions. The authors acknowledge the core-facilities and technical assistance of the NCBES qPCR Facility, Centre for Microscopy and Imaging (CMI), Flow Cytometry Core Facility, funded by NUI Galway and NDP 2007-2013, Cycles 4 and 5. The authors also acknowledge the support of NANOREMEDIES, which is funded by the Programme for Research in Third Level Institutions, Cycle 5 and co-funded by the ERDF. The AFM used for this research work was funded by Science Foundation Ireland (SFI07/IN1/B931).

We thank Prof. Peter Dockery and Dr. Kerry Thompson for their advice and technical assistance from the Anatomy Department. We would also like to thank Mr. Maciek Doczyk for the illustrations in the manuscript.

