## Supplementary Information for "Temporal Changes Guided by Mesenchymal Stem Cells on a 3D Microgel Platform Enhances Angiogenesis In Vivo at a Low-Cell Dose"

### SI Materials and Methods

#### Culture of Cells

hMSC primary cells were isolated from fresh human bone marrow. Cells were isolated in the same media as used during the passaging, so the cells were never exposed to more than one type of serum. Non-adherent cells were removed after three days, after which the media was changed twice per week. Isolated cells were characterised based on the routine tri-lineage differentiation potential and the presence of cell surface markers using flow cytometry. hMSCs were cultured in complete medium (MEM alpha, GlutaMAX™ supplemented with 10% foetal bovine serum and 1% penicillin/streptomycin) and maintained at 37°C in a humidified atmosphere containing 5% $CO_2$. For both *in vitro* and *in vivo* experiments MSCs were pooled from four to six unrelated bone marrow donors, and after routine characterisation and were frozen down at passage P1.

#### Assessment of Cell Morphology

Microgels were imaged for four days' post-fabrication for changes in cell morphology. Microgels were briefly washed in PBS and incubated with 10mM calcein AM (Fluka, Germany) for ten minutes. Samples were then imaged as Z-Stacks through 500-800μm on an Andor™ Olympus Spinning Disk Microscope (Andor, Belfast, Northern Ireland), using Andor™ IQ software, with an 20X oil immersion objective lens. Green viable cells were visualised by excitation with the 488nm laser line. Sample images were obtained as Z-slices 5μm apart. Each Z-stack was analysed using the Volocity® 5.0 software (Perkin Elmer Inc., Waltham, USA). Cell morphology parameters such as shape factor index, surface area-to-volume ratio and longest cell axis distance were analysed. Shape factor is the ratio of surface area of a sphere to the surface area of the object. In the equation below $A_0$ is the surface area and $V_0$ is the volume.

$$\text{3D Shape Factor} = \frac{\pi^{\frac{1}{3}}(6V_0)^{\frac{2}{3}}}{A_0}$$

Datasets were analysed using defined algorithms to estimate a series of measurements related to the dynamic changes in cell morphology across three different planes (top, middle and bottom) of the microgels.

#### Assessment of Changes in Microgel Modulus Seeded with hMSCs

For measuring the Young's modulus of the cell seeded microgels incubated for 24 or 96 hours, atomic force microscopy was utilised. All the measurements were performed with MFP-3D-BIO AFM (Asylum Research/Oxford Instruments, US) fitted on a Ti/E inverted microscope (Nikon, Japan) and run with Igor Pro™ 6.34A software (WaveMetrics, US) with Asylum Research plugin (ver. 120804+2209). Probes were prepared by fixing 10μm silica glass spheres (Windsor Scientific, UK) at the end of type B CSC38 tipless AFM cantilevers (Mikromasch, Bulgaria) with epoxy glue. Stiffness and sensitivity of the cantilevers were calibrated in air (Sader, non-contact method) and subsequently in PBS (contact method) to correct for changes in lever sensitivity. These values ranged between 62.25 and 83.25nm/V for sensitivity (in liquid), and between 64.7 and 73.15pN/nm for lever spring constant.

Microgels suspended in PBS were deposited on a microscope glass slide and allowed to settle down for ten minutes. This sample preparation appeared to prevent the microgel from rolling during indentation; however, movement of the microgel or rolling of the spherical tip on the surface could lead to errors in quantification of the modulus. For each condition, namely microgels prepared with 1, 2 or 3mg/ml collagen which were seeded with cells (following 24 or 96 hours incubation and fixing with alcohol), 3-4 microgels were chosen and 36-68 measurements were performed per group (4-29 per microgel). Each measurement consisted of an indentation with constant a loading force of 100pN, 3-5μm force distance and 1μm/s tip approach/retraction speed. Used force was optimised to avoid indentation depths greater than 3μm (30% of the silica sphere tip) on the softest sample. The measurements on the microgels were performed in liquid, and the resulting force-distance curves were processed with Asylum plugin for Igor using a Hertz model (1).

Indenter (silica sphere) modulus was assumed as 68.00GPa and its Poisson's ratio was assumed as 0.19. Considering the viscoelastic properties of a swollen hydrogel, its Poisson's ratio was set to 0.5 (2). The whole curve length was used for fitting, except in the cases where good fit quality could only be achieved by fitting only the contact portion. Fit quality was assessed with a reduced chi square value ($\chi^2$) calculated for each modulus and was further used to obtain a weighted average modulus for a given sample type. The closer $\chi^2$ was to 1, the better the fit.

**Analysis of Cell Viability**

Microgels were then digested in collagenases for 20 minutes and sterile filtered with 70μm filter mesh to harvest cells. Before analysis on BD FACS Canto (BD Biosciences, USA), cells were re-suspended in propidium iodide/RNase staining buffer (BD Biosciences, Ireland). Live cells (PI-negative) and dead cells (PI-positive) cells were included for analysis using FlowJo software 8.5.2 (Tree Star, Ashland, OR).

**Scanning Electron Microscopy (SEM) Imaging of hMSC Embedded Microgel**

Microgels were washed with phosphate-buffered saline (PBS) twice and fixed in 2.5% glutaraldehyde for three hours. Then, the samples were dehydrated in a graded ethanol series and air-dried. After gold sputtering, the microgels were examined with a Hitachi S-4700 scanning electron microscope (Hitachi Scientific Ltd., Japan).

**Ultrastructure Analysis of Cellular Inclusions**

Microgels were fixed for four hours at room temperature in 2.5% glutaraldehyde at pH 7.4, post-fixed in 1% osmium tetroxide in 0.2M sodium cacodylate buffer at pH 7.4 for one hour. The microgels were dehydrated in a series of graded ethanols and infiltrated with a mixture of propylene oxide and epoxy resin (Agar Low Viscosity Resin kit, Agar Scientific, UK), and finally infiltrated with 100% resin on a rocker overnight. The microgel embedded resin blocks were then cured in an oven at 65°C for 48 hours. Ultrathin sections were cut and transferred to standard transmission electron microscope (TEM) grids and contrasted with uranyl acetate before being viewed under TEM for imaging. TEM images of the cell embedded microgels were transformed to an 8-bit image (Fiji-ImageJ software) (3). Subsequently, a classifier plugin was trained on the TEM micrographs using Trainable Weka Segmentation plugin, defining dark cellular inclusions and all other intracellular organelles (background). The total cell area and the number of cellular inclusions were quantified using automated threshold of the respective probability map image and object counter analysis of Fiji/ImageJ with an average inclusion size less than 500$\mu m^2$.

**Cell Surface Marker Analysis of hMSCs**

Flow cytometry analysis was used for confirming the presence of human mesenchymal stem cell surface markers. The cells were harvested by collagenase digestion of microgels and washed twice in PBS. The cells were re-suspended in FACS buffer (PBS, 2% FBS and 0.1% NaN3). Approximately, 5-7 × $10^4$ cells were incubated with anti-human primary monoclonal antibodies CD90, CD105, CD73, CD45, CD34, CD14, and CD20 (Miltenyi Biotec, Bergisch-Gladbach, Germany). Data was acquired by BD FACS Canto (BD Biosciences, San Jose, CA) FACS Calibur flow cytometer and analysed by FlowJo software (TreeStar Inc., OR, USA).

**Immunostaining for $\alpha V\beta_3$ and Vitronectin**

Microgels cultured for 96 hours were fixed in 4% paraformaldehyde for 30 minutes and blocked with 10% normal goat serum for two hours. Microgels were then incubated with mouse anti-human $\alpha V\beta_3$ (Santa Cruz Biotechnology) and rabbit anti-human Vitronectin (Santa Cruz Biotechnology) primary antibody (1:200) in 2% goat serum overnight at 4°C. Alexafluor® 488, goat anti-mouse IgG (1:100) and Alexafluor® 594, goat anti-rabbit IgG (1:100) secondary antibody was incubated for one hour at room temperature in blocking buffer. Following six PBS washes, nuclei were counterstained with Hoechst (Life Technologies) for five minutes. Samples were then imaged as Z-Stacks on Andor™ Olympus Spinning Disk Microscope (Andor, Belfast, Northern Ireland), using Andor™ IQ software, with an X 20 objective lens. Calculation of protein expression and Pearson's co-localisation coefficients were calculated in Volocity® 5.0 software (Perkin Elmer Inc., Waltham).

**Histology and Immunohistochemistry**

Three weeks after the treatment mice were sacrificed to harvest the tissue. Hindlimb gastrocnemius and lower quadriceps muscles were carefully dissected and weighed before being fixed in 4% PFA for 48 hours. The samples were then rinsed with PBS before being snap frozen followed by embedding in Optimal Cutting Temperature (OCT) compound (TISSUE-TEK®; Sakura Finetek USA. Inc.). Cryosections were taken at a thickness of 7µm from four different depths at 50µm intervals using a Leica CM1850 cryostat (Leica Microsystems, Germany) set at -22°C. The samples were stained with hematoxylin and eosin (H&E). For immunofluorescence, cryosections were treated with 1X proteinase K solution and incubated at 37°C for 20 minutes for antigen retrieval. The tissue sections were blocked with 1% BSA for two hours to avoid non-specific binding. Anti-mouse polyclonal primary antibody specific for CD31/PECAM-1 (Abcam, Ireland) endothelial marker or CD68 (Abcam, Ireland) macrophage marker were incubated overnight at 4°C. Secondary antibody labelled with AlexaFluor® 488 or AlexaFluor® 594 (1:500, Invitrogen, Ireland) was applied for one hour at room temperature, followed by nuclear counter staining using Hoechst dye (Life Technologies).

**Lectin Histochemistry**

For lectin histochemistry, the muscle sections were washed two times with distilled water for five minutes each. Slides were washed with Tris-Buffered Saline (TBS) 1X enriched with $Ca2^+$ and $Mg2^+$ cations (20 mM Tris-HCl, 100 mM NaCl, 1 mM CaCl2, 1 mM MgCl2, pH 7.2) and then blocked with 2% periodate-treated bovine serum albumin in TBS for one hour. Sections were then incubated with either Wheat germ agglutinin (WGA) (10µg/ml) or Ulex Europaeus Agglutinin I (UEA-1) (15µg/ml) lectin labelled with fluorescein isothiocyanate (FITC) (EY Labs Inc.) in TBS for one hour. The sections were washed three times with TBS and nucleus was counterstained with DAPI (1:10000). All slides were cured in dark overnight prior to imaging on an Andor™ Olympus Spinning Disk Microscope (Andor, Belfast, Northern Ireland).

**Model of Hindlimb Ischemia**

All the animal treatment groups and experimental procedures were approved by the institutional Ethics Committee. A recently developed double ligation hindlimb ischemia nude mouse model (4)was used to test the therapeutic efficacy of the optimised 2mg $ml^{-1}$ and 0.8 million cells microgel construct. The mice were anesthetized with intra-peritoneal injections of xylazine (10 mg⁄kg) and ketamine (80 mg⁄kg). For hindlimb ischemia induction, two separate skin incisions were made; one around the groin area and the other above the knee. The artery was then carefully separated from the vein and a ligation was placed before the profunda femoris. This was followed by a second ligation placed before the saphenous–popliteal bifurcation branch. A cut was then made between the ligation sites interrupting the blood flow to the limb. Mice were divided into the following treatment groups (n=12/group): hMSC embedded microgels (30±5, 1600 cells/microgel), $5x10^4$ cells (low-cell density; approximately the same number of cells embedded in microgels), $1x10^6$ cells (high-cell density), microgels alone and saline which were implanted or injected at the proximal artery ligation site after surgical induction. The incision was then sutured closed and the animals were given analgesic of 0.05mg/kg buprenorphine prior to recovery from anaesthesia. Laser Doppler measurement was acquired weekly for three weeks, along with the assessment for improvement in ambulation and signs of necrosis or auto-amputation. The animals were sacrificed at the end of the three weeks to harvest the tissue.

**Assessment of Ambulation and Necrosis**

The efficacy of the treatment was evaluated semi-quantitatively by gross examination of the foot. Limb condition was classified into several levels of severity and scored based on signs of necrosis and the extent of plantar flexion or dragging observed on the operated foot while walking. A point-based scale was devised to quantify the extent of necrosis in the limb with 0=blue correlated with no necrosis to 5=black indicative of severe necrosis. Similarly, ambulatory impairment was graded between 0 and 3, where 0 refers to normal ambulatory function similar to the non-operated foot and 3 indicates poor ambulation manifested by dragging of the foot. Two blinded independent scorers carried out the scoring every week for three weeks.

**Histomorphometry**

Histomorphometric analysis is typically performed to quantify tissue responses to injectable or implantable therapeutics. It allows assessment of angiogenesis in vascular beds and infiltration of inflammatory cells(5). In order to avoid observational bias in sampling, a systematic methodology was employed throughout the study as previously described(5-9). For quantification of the blood vessels and infiltration of inflammatory cells, twenty images per sample per animal were analysed across four levels separated by 50μm. The 'forbidden line' method was used to count the blood vessels that run parallel depending on how the vessel intersects or does not intersect in the counting frame(10). This quantification is robust and highly representative of the changes that occur in tissues. In the current study 70-80 tissue sections per animal in each group were analyzed.

The fields were captured at 40X magnification. A grid mask with squares of defined dimensions was placed on the histology sections using Image Pro® Plus software. Quantified parameters were capillary density, radial diffusion and the area fraction of infiltrating inflammatory cells.

*Vessel Density and Radial Diffusion*

For angiogenesis, the points where the squares on the grid intersected blood vessels were numbered on the section. The surface capillary density of blood vessels was calculated using the intersected blood vessels multiplied by the area of the total area of the tissue within the square. Radial diffusion was calculated using the formula (1/SQRT($L_v$*PI)*1000, which is the distance between blood vessels and is an indicator of the capillary network in the vascular bed. The shorter the distance between blood vessels, the smaller the distance required for nutrients to diffuse into surrounding tissues.

*Area Fraction of Inflammatory Cells*

The area fraction of infiltrating inflammatory cells was also evaluated. Inflammatory cells were counted and these cells included lymphocytes and neutrophils. The area fraction was determined with a 1938 points grid using Image Pro® Plus software (Media Cybernetics). Neutrophils were identified as small dense circular multi-lobed cells and lymphocytes as small round dense cells with large nuclei.

**Tissue mRNA Extraction and Mouse Endothelial PCR Arrays**

RNA from muscle tissues was stored in RNAlater® (Thermo Fisher Scientific, UK) and extracted using RNAeasy® Microarray Tissue mini kit (Qiagen GmbH, Germany). Tissues were homogenised using TissueLyser LT(Qiagen) for 20 minutes at 50Hz. Extracted RNA samples were treated with DNase to remove contaminating genomic DNA. RNA quality was assessed using the Agilent 2100 Bioanalyzer (Agilent Technologies, CA) and samples with RNA Intergrity Number (RIN) RIN >7 were used for downstream cDNA conversion with a $RT^2$ first-strand kit (Qiagen GmbH, Germany) according to the manufacturer's protocol. Eighty-four gene targets were analysed on a Mouse Endothelial Cell Biology $RT^2$ profiler PCR array (PAMM-015Z, SaBiosciences Corp., USA). Statistical analysis of relative gene expression results in real-time PCR was performed using REST© software tool (11). Briefly, a Pair-Wise Fixed Reallocation Randomization Test was used to determine significant differences between the treatment and the control groups (n=3/treatment group; n=1 pooled tissue samples from four animals). A p-value less than 0.05 was considered to be significant.

**Ingenuity Pathway Analysis**

The genes for *in vitro* and *in vivo* PCR-arrays were uploaded to Ingenuity Pathway Analysis (IPA) software. Pathway analysis was performed from information contained in the IPA knowledge base (IPKB). The database was limited to human, mouse and rat species. For network analysis, IPA computed a score (p-score=-log10(p-value)) according to the fit of the set of supplied genes and a list of biological functions in IPA. The score considers the number of genes in the network and the size of the network to approximate how relevant this network is to the original list of genes and allows the networks to be prioritized for further studies. A z score ≥2 (or ≤−2) ($p<0.05$) was considered significantly activated or inhibited.

**Tissue Sample Processing and MALDI Mass Spectrometry Imaging for *N*-Glycans**

Unfixed muscle tissue samples from the upper gastrocnemius were stored at -80°C for mass spectrometry imaging (MSI) and MALDI-TOF analysis after in-solution glycan release. For MSI experiments and H&E staining, consecutive cryosections of 15μm in OCT were cut using Leica CM3050 (Leica Microsystems, Germany) and thaw-mounted on conductive indium tin oxide (ITO)-coated glass slides (Delta technologies, USA). Frozen tissues were thawed and dried in a vacuum desiccator for one hour at room temperature before washing. Tissue sections were washed in double distilled water ($ddH_2O$) and sequentially dehydrated in graded ethanol solution (50-100%) for 30 seconds each. Dehydrated sections were washed in $NH_4HCO_3$ (10mM, pH 8.0) twice for five minutes and air dried at room temperature. Glycerol-free PNGase F (200 μl) (New England Biolabs, UK) was dialysed in $NH_4HCO_3$ (25mM, pH 8.0) and applied on tissue (15nL at 100 μm spacing) using a ChIP-1000 (Shimadzu, Japan). No-enzyme controls were printed using the $NH_4HCO_3$ buffer. Tissue sections were incubated overnight at 37°C in a humidified chamber, and a defined peptide mix was spotted adjacent to the section for TOF calibration. 2,5-Dihydroxybenzoic acid (DHB) matrix (5.5 mg/mL) in 0.1% TFA containing 1 mM NaCl was deposited on the specimens using a home-built sublimation apparatus (Marchetti-Deschmann laboratory, TU Wien), and recrystallization of the matrix was performed by hydrating the slides in vapours of acetic acid in a humidified chamber (12). MSI data was acquired on a MALDI TOF/TOF (ultrafleXtreme™, Bruker Daltonics, Germany) mass spectrometer in reflectron mode (using a spatial resolution of 50 μm and 50 shots per position of a random walk within each pixel) with glycan masses ranging up to *m/z* of 2500. Flex Imaging™ v3.0 (Bruker Daltonics, Germany) was used for data and image processing. All presented images are total ion current (TIC) normalized.

**In-solution Tissue Digest for *N*-Glycan Release**

Muscle tissue chunks from the upper gastrocnemius were microdissected and placed in eppendorfs for washing. Tissues were washed once with ultra-high-quality water ($ddH_2O$) with an additional two washes (2 x 5 minutes) with 10mM $NH_4HCO_3$. The tissues were incubated in 10mM $NH_4HCO_3$ at 95°C for 30 minutes. Further, tissues were gently homogenised with tweezers in the eppendorf tube and centrifuged at 8,000RPM for ten minutes. The supernatant tissue lysate was incubated with glycerol-free PNGase F at 37°C overnight. The mixture was deposited onto a ZipTip C18 tip, and glycans (together with unbound proteins/peptides) were obtained in the flow through and spotted with DHB matrix onto a MTP-384 AnchorChip™ target (Bruker Daltonics) and allowed to dry at room temperature. For calibration, a standard peptide mix was spotted separately onto calibrant AnchorChip spots. MS data was obtained on the same instrument described above using FlexControl™ v.3.4 (Bruker Daltonics). For analysis, automatic peak picking was performed using a signal-to-noise (*S/N*) threshold of 7. Peak identity and intensities of *m/z* values were matched with MSI experiments for the same sample and represented as ion intensity maps on the tissues. The target parent ions were fragmented using the LIFT mode and the resultant MS/MS spectra was screened for possible peptides using MASCOT (http://www.matrixscience.com/) and de novo sequencing approaches (manual assignment of mass differences). Ions not identified as peptides were submitted as $[M+Na]^+$ and with a mass accuracy of ±2 Da to GlycoMod (http://us.expasy.org/tools/glycomod) to identify possible N-glycan structures (phosphate, sulphate modifications and presence of sialic acids excluded).

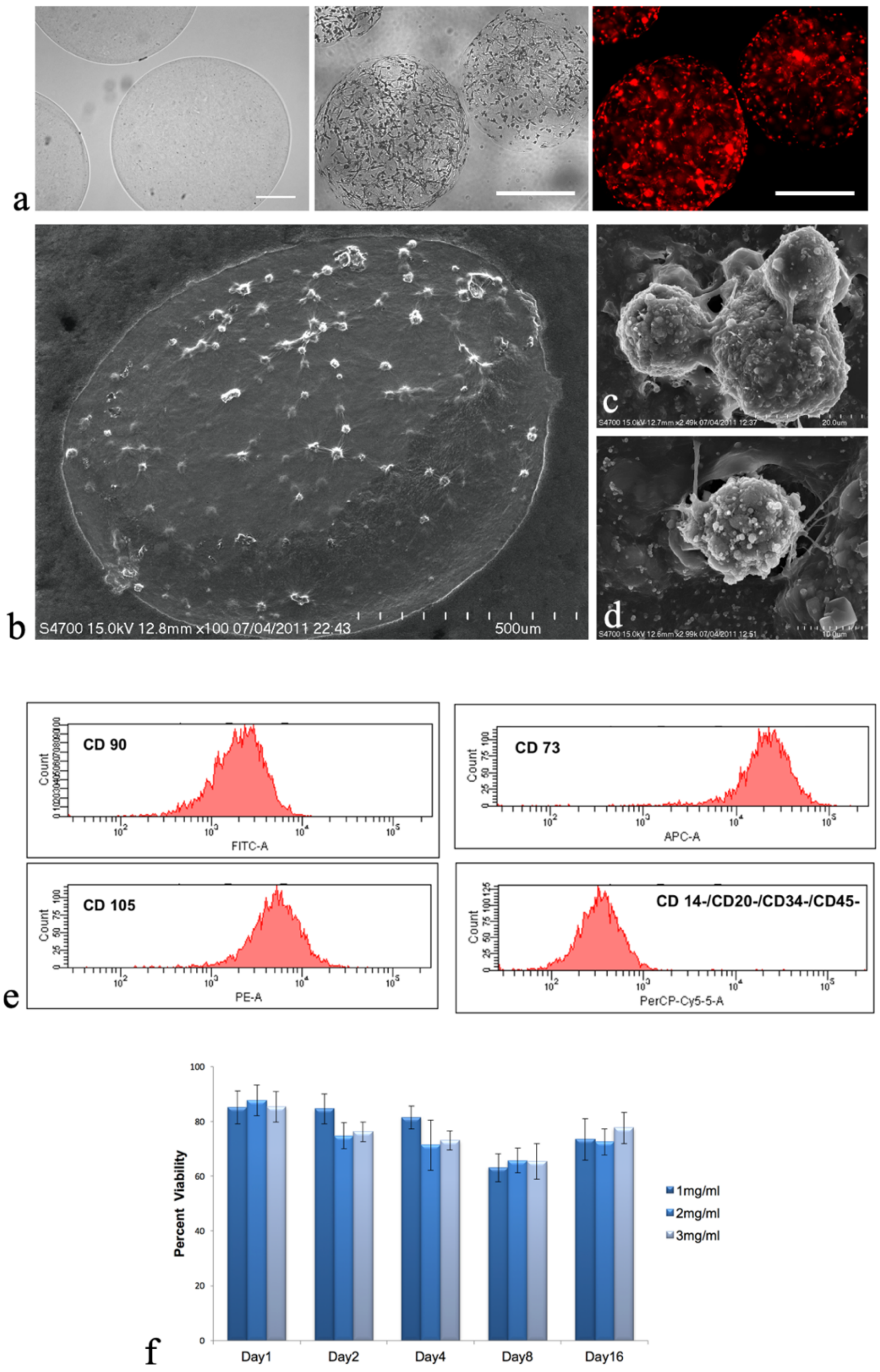


**Fig. S1: (a)** Microgels unseeded (left); scale bar 200μm, hMSC embedded (center) and PkH-26 labelled hMSCs (right) embedded in microgels after 48hours in vitro (scale bar 500μm).**(b)** SEM micrographs showing topographical appearance of a hMSC seeded microgel with **(c)** cell-cell and **(d)** cell-matrix interaction **(e)** Flow cytometry analysis showing retention of phenotypic human mesenchymal stem cell surface markers 14 days post-seeding on microgels. **(f)** Over 80% cell viability observed for 16 days in all the microgel groups (collagen concentrations 1mg ml-1, 2mg ml-1 and 3mg ml-1) at 0.8 million cells ml-1 cell density. n=6, $p<0.05$.

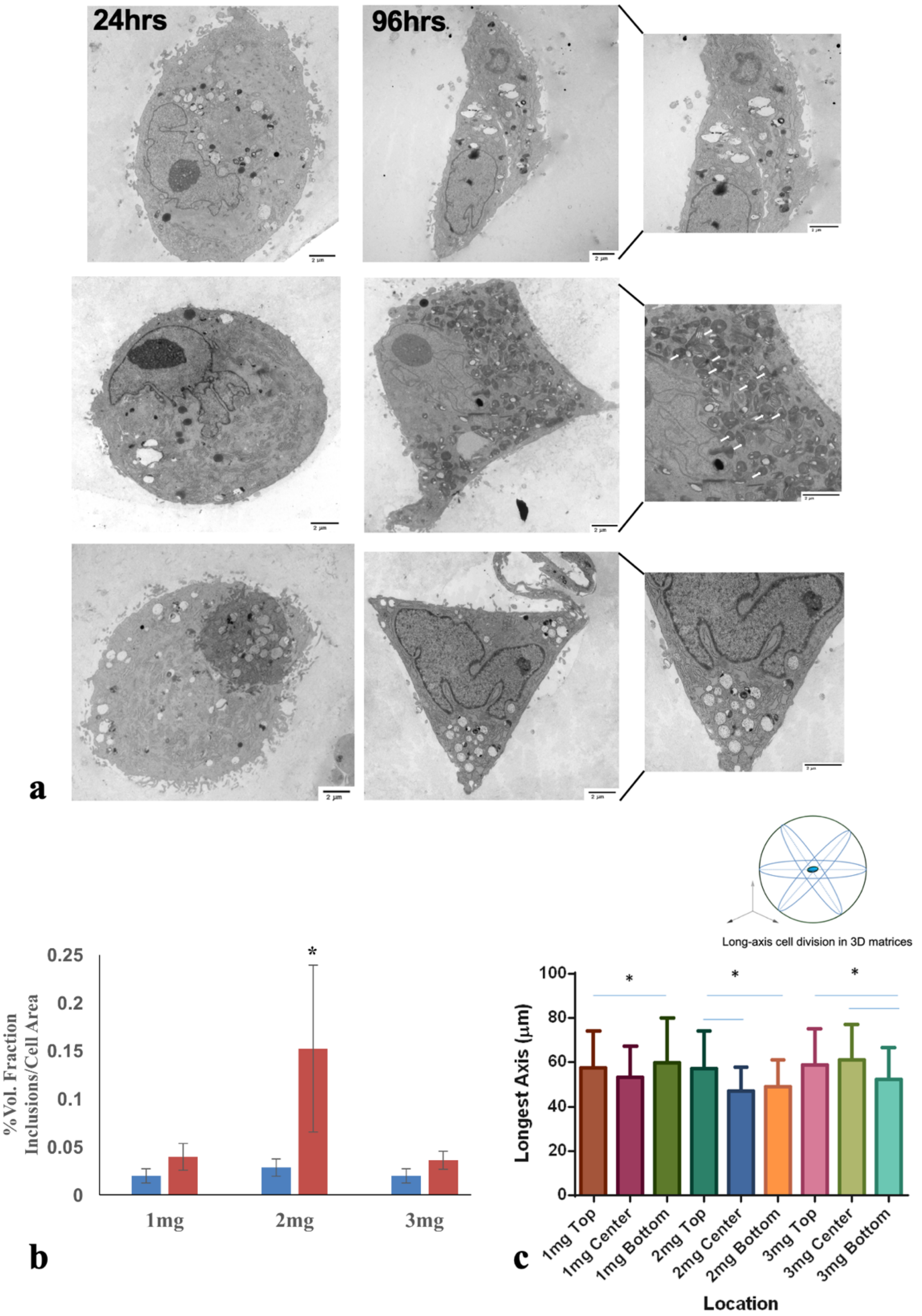


**Fig. S2:** Preconditioning results in accumulation of intracellular inclusions in hMSCs embedded in microgels. **(a)**TEM micrographs of hMSCs cultured for 96 hrs containing darkly stained cellular inclusions (indicated with arrows) with distinct cell morphology. Scale bar, 1 μm. **(b)** Quantification of intracellular inclusions in 1, 2 and 3 mg $ml^{-1}$microgels (n=5) using FIJI's Trainable WEKA segmentation plugin. **(c)** Differences in hMSC alignment on its longest axis on collagen microgels indicative of cellular anisotropy. *indicates statistical significance ($p<0.05$).

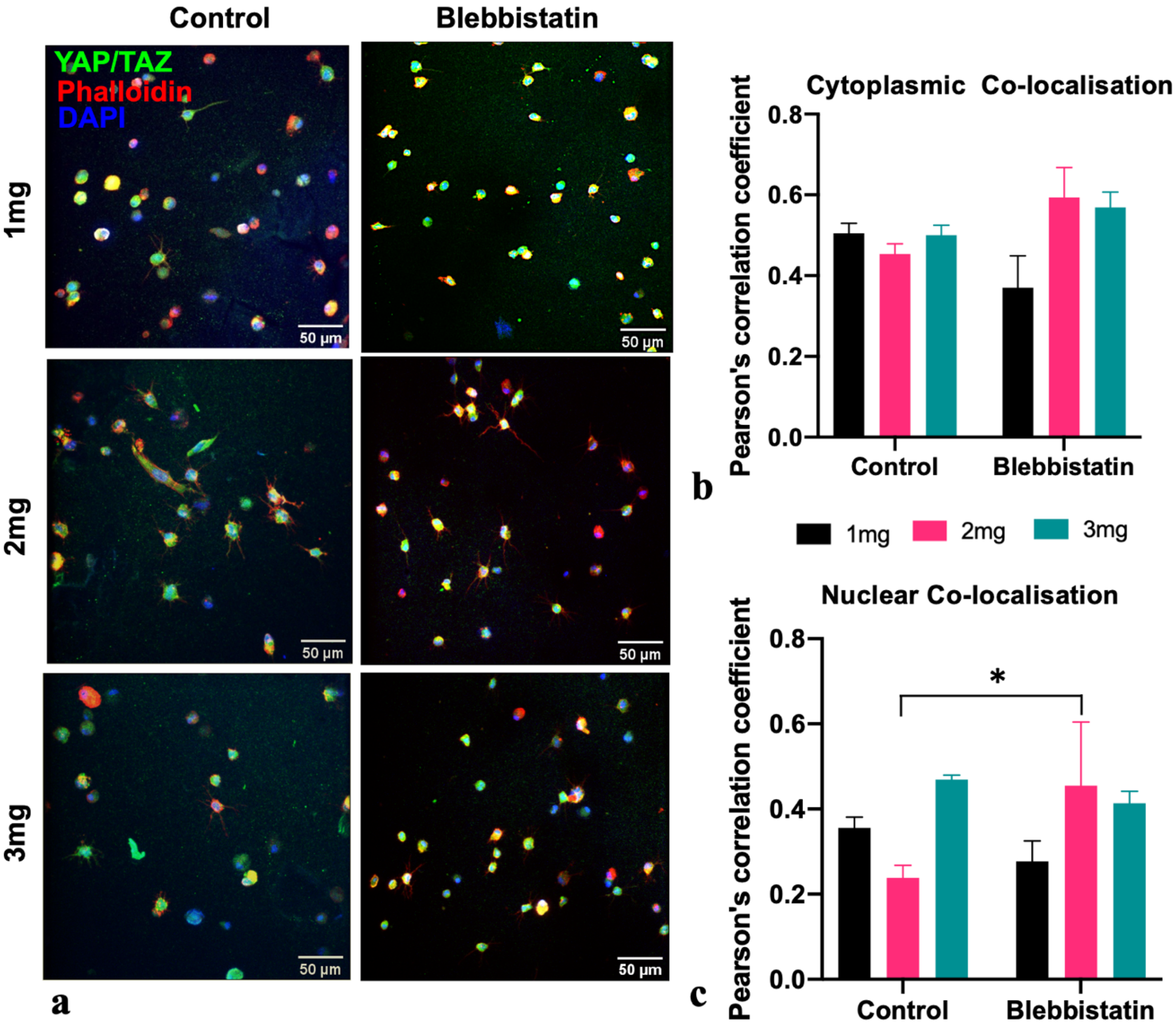


**Fig. S3**. **(a)** Immunostaining of YAP/TAZ (green), cytoskeleton (red), and nucleus (blue) of human mesenchymal stem cells in 1, 2 and 3mg $ml^{-1}$ microgels at 96 hrs pre-treated with myosin inhibitor blebbistatin. Magnification 20X, Scale bar, 50µm. **(b)** Quantification of cytoplasmic and nuclear YAP/TAZ co-localised with the cytoplasm (Phalloidin) or nucleus (DAPI) using Pearson's coefficient for co-localisation. Significant increase in nuclear YAP/TAZ co-localisation observed in 2mg $ml^{-1}$ (n=4) blebbistatin group compared to 2mg $ml^{-1}$ control. *indicates statistical significance ($p<0.05$).

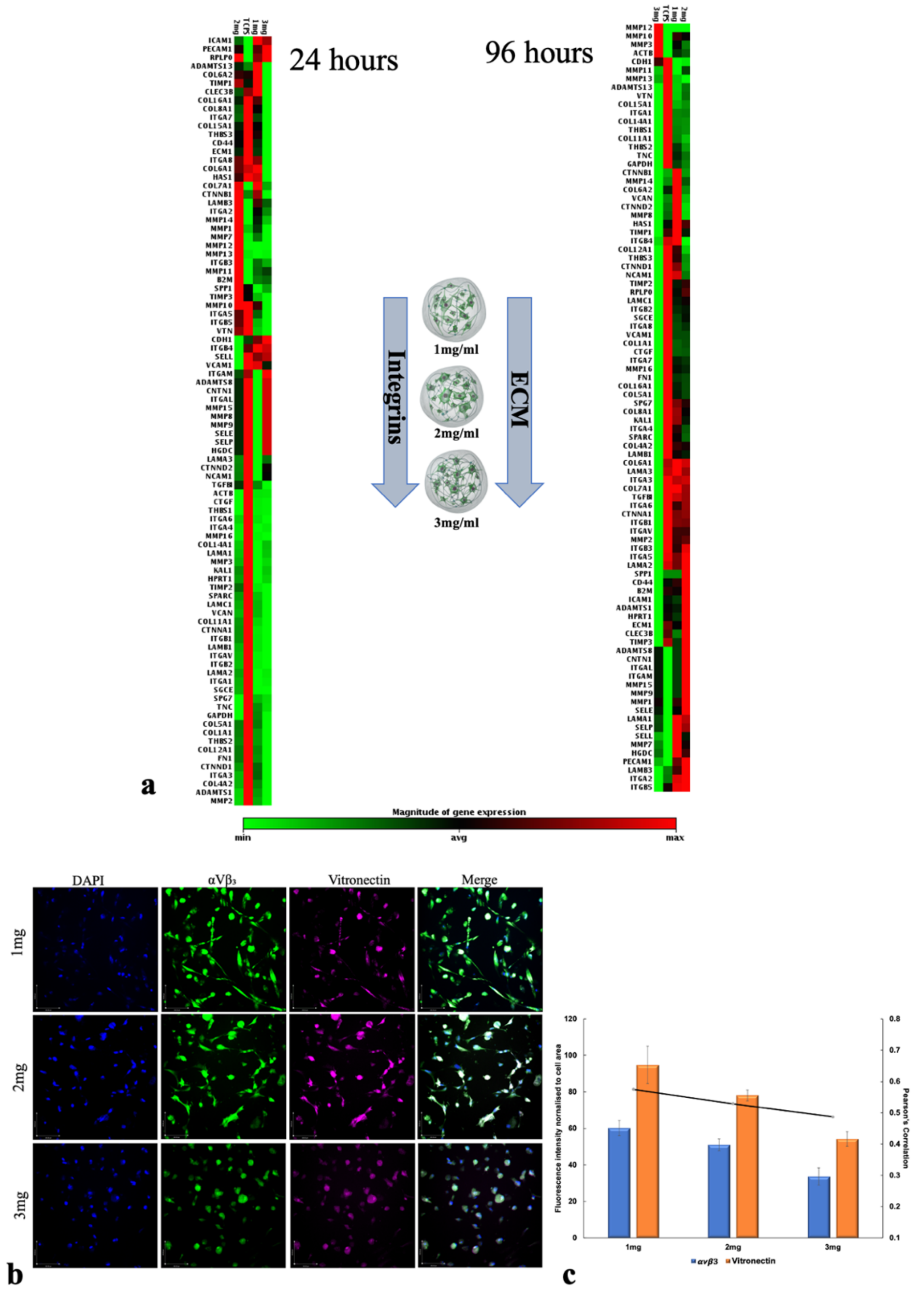


**Fig. S4: (a)** Heatmap of extracellular matrix and cell adhesion gene markers showing differential fold expression and microgel concentration (1mg $ml^{-1}$, 2mg $ml^{-1}$ and 3mg $ml^{-1}$) dependent modulation of genes compared to plastic adherent human mesenchymal stem cells at 24 and 96 hours. **(b)** Immunostaining of integrin $\alpha V\beta_3$ (green), vitronectin binding domain (magenta), and nucleus (blue) of human mesenchymal stem cells in 1, 2 and 3mg $ml^{-1}$ microgels at 96 hrs. Magnification 20X, Scale bar, 190µm. **(c)** Quantification of protein expression using fluorescence intensity and Pearson's coefficient for co-localisation. A linear reduction in expression of integrin $\alpha V\beta_3$ and vitronectin is observed with the increase in microgel concentration at 96 hrs (n=3). *indicates statistical significance ($p<0.05$).

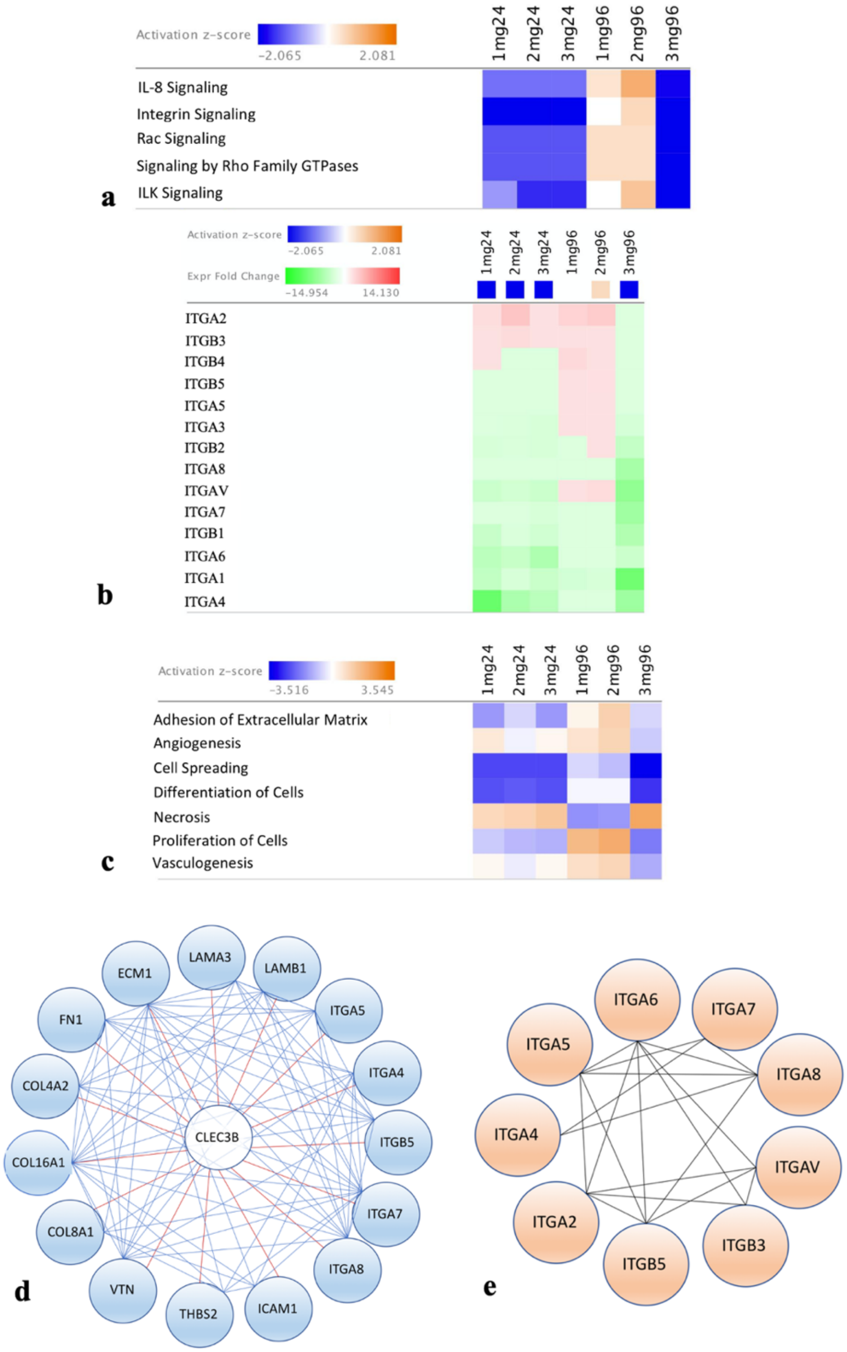


**Fig. S5: (a)** Heatmap of *z*-scores (indicating activation or inhibition) of canonical pathways Ingenuity Pathway Analysis (IPA) with a threshold of 1.3-fold expression, $p<0.05$. The dataset represents changes in gene expression at 24 hrs and 96 hrs. **(b)** Heatmap of genes in the integrin signalling network expressed as expression fold change **(c)** Heat map displaying *z*-scores of biological functions (IPA).**(d)** Pearson's correlation analysis confirm strong (Pearson's $p>0.6$) and significant ($p$ value $<0.05$) correlations of gene markers to CLEC3B, identified as the highly enriched node in 2mg $ml^{-1}$ microgel condition at 96 hrs. **(e)** Strong (Pearson's $p>0.6$) and significant ($p$ value $<0.05$) correlation between integrins in 2mg $ml^{-1}$ microgel condition at 96 hrs.

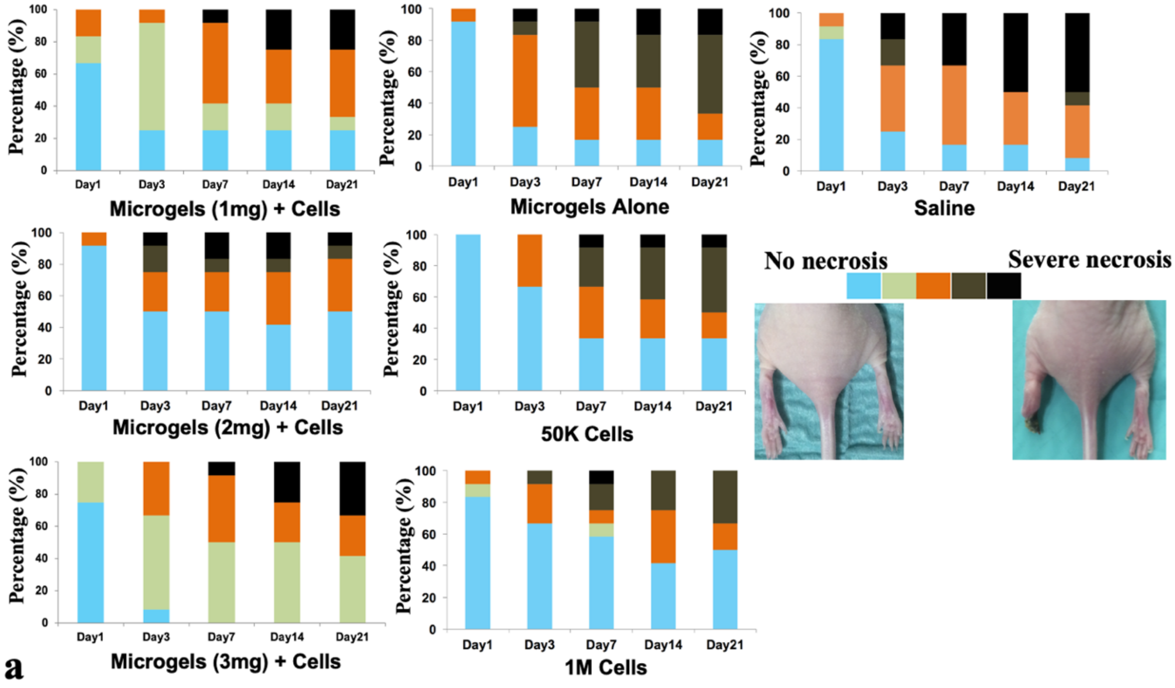


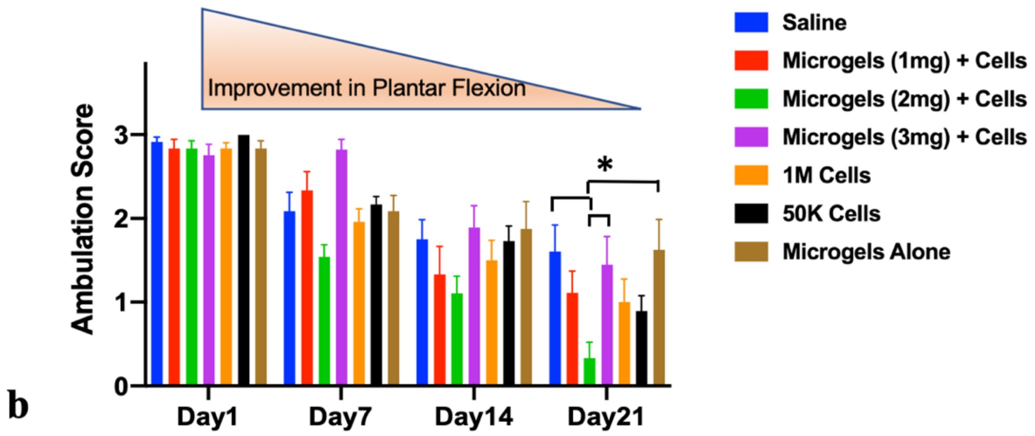


**Fig. S6**. **(a)** Higher limb salvage in animals treated with hMSC embedded in microgels at a low-cell dose. At day 21 treatment group microgels with hMSCs showed higher limb salvage with 50% (6 out of 12) of the mice protected from tissue damage or necrosis and 15% showing varying degree of moderate to severe necrosis (n=12/group). The color-coded scale is based on a 5-point modified Tarlov scale; 0= Normal, 1= mild redness or cyanosis of toe tips, 2= cyanosis of toes with mild necrosis of toes, 3= moderate necrosis (affecting two or more toes), 4= Severe necrosis affecting the metatarsals, 5= autoamputation of toes/distal limb. **(b)** Significant improvement in ambulation at day 21 in mice treated with hMSC embedded in microgels at a low-cell dose (n=12, $p<0.05$). Hindlimb scoring (index of muscle function); 0= normal, 1= plantar flexion but no flexion of toes, 2= no plantar flexion or dragging and 3= dragging of the foot. A higher score indicates impaired ambulatory function.

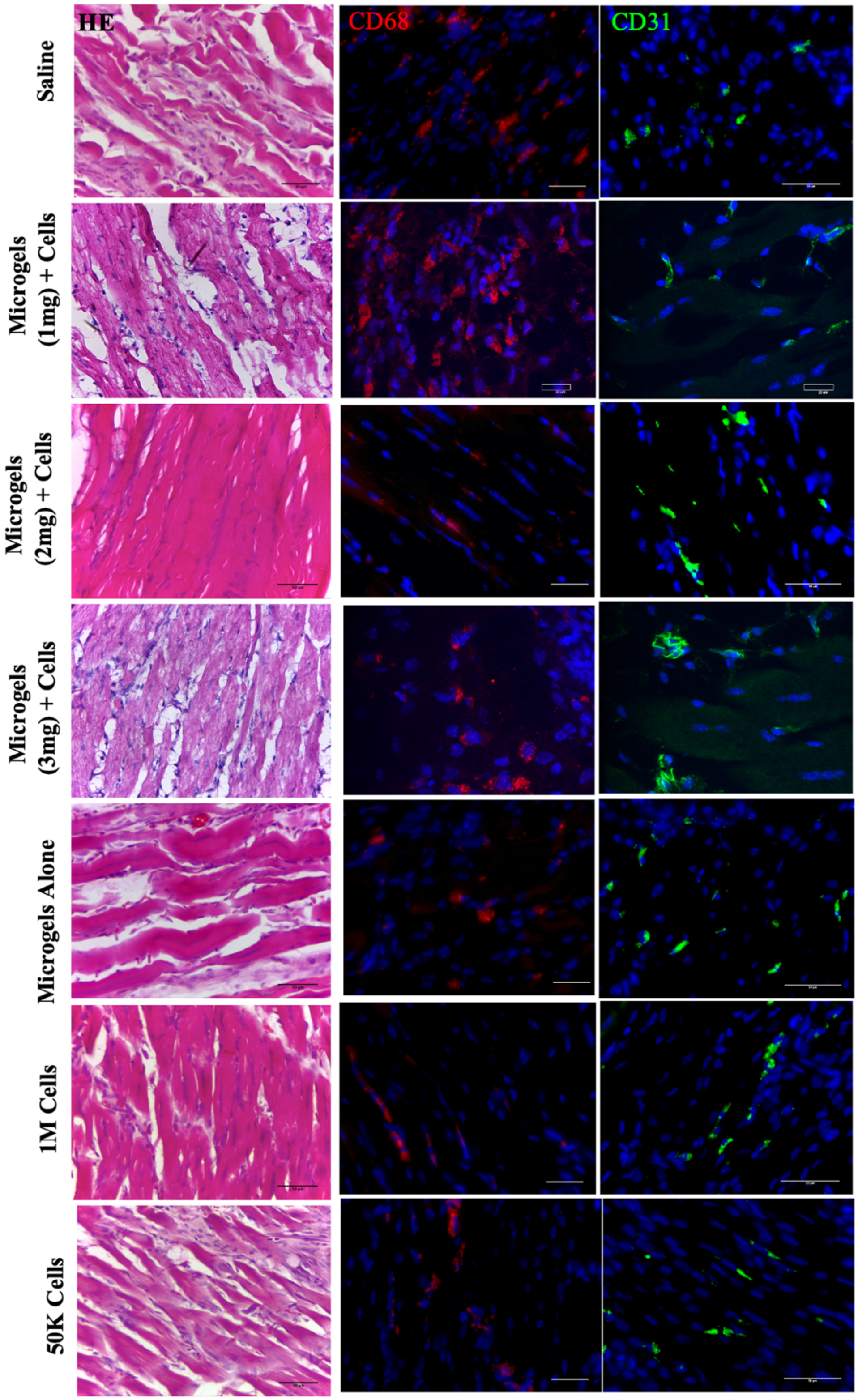


**Fig. S7**. Immunohistochemical and H &E staining of muscle tissues from treatment groups. CD68 (red) staining for macrophages and CD31 (green) staining for blood vessels.

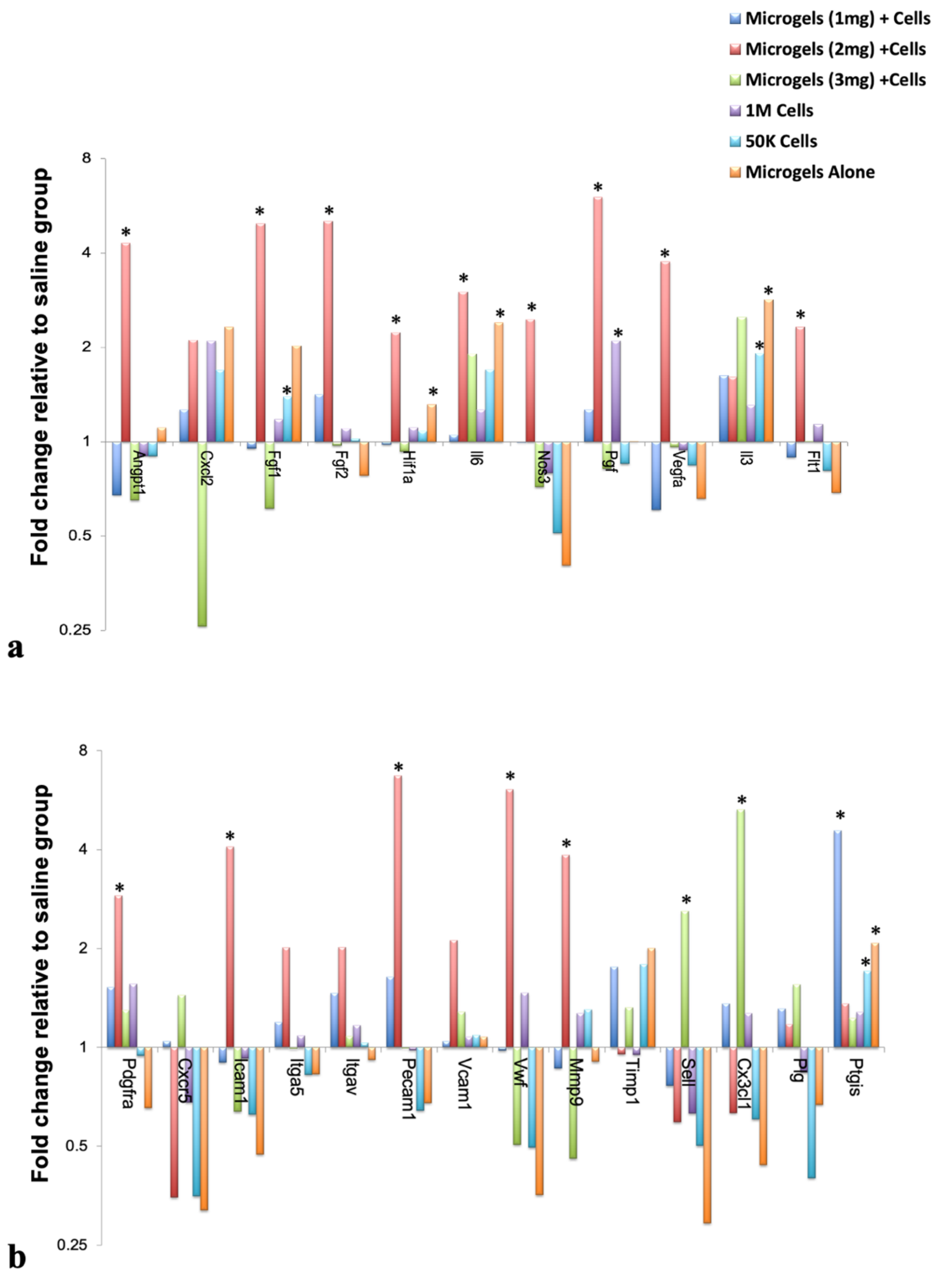


**Fig. S8**. Significantly high upregulation of genes associated with angiogenesis **(a)** Growth factors and cytokines and **(b)** Extracellular matrix and cell surface markers were observed in mice treated with microgels with hMSCs post-operative at day 21. Statistical significance tested with two-way ANOVA, $p<0.05$; (n=3/group; n=1 pooled from 4 animals). *indicates statistical significance ($p<0.05$).

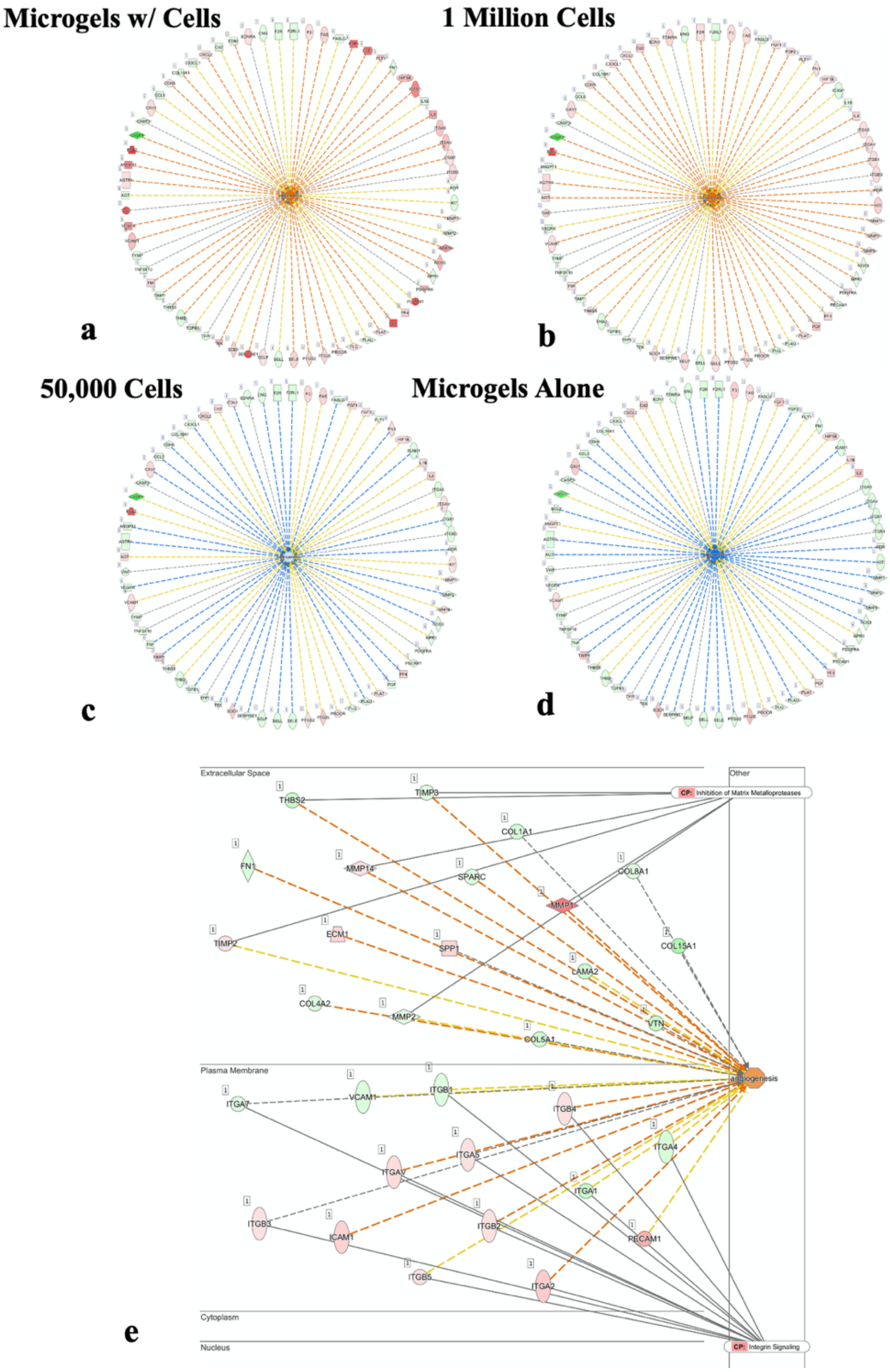


**Fig. S9**. Ingenuity Pathway Analysis predicted a downstream effect with increased angiogenesis *in vivo* treated with **(a)** microgels with cells (Z-score 2.53) compared to **(b)** 1 million cells (Z-score 2.34), **(c)** 50,000 cells (Z-score -0.24) and **(d)** microgels alone (Z-score -1.78). In microgels with cells group 40 of 68 genes had an expression direction consistent with increased angiogenesis, yielding a Z-score of 2.53. Red symbols indicate increased transcript levels, while green symbol indicate reduced transcript levels. The dotted lines indicate indirect relationships leading to activation (orange) or inhibition (blue). Yellow and grey dotted lines indicate inconsistent relationships and no predicted effects, respectively. **(e)** Molecular network induced by hMSCs embedded in 2mg ml$^{-1}$ microgels. Upregulated (red) and downregulated (green) genes resulting in the activation of angiogenesis pathway as revealed by IPA software analysis.

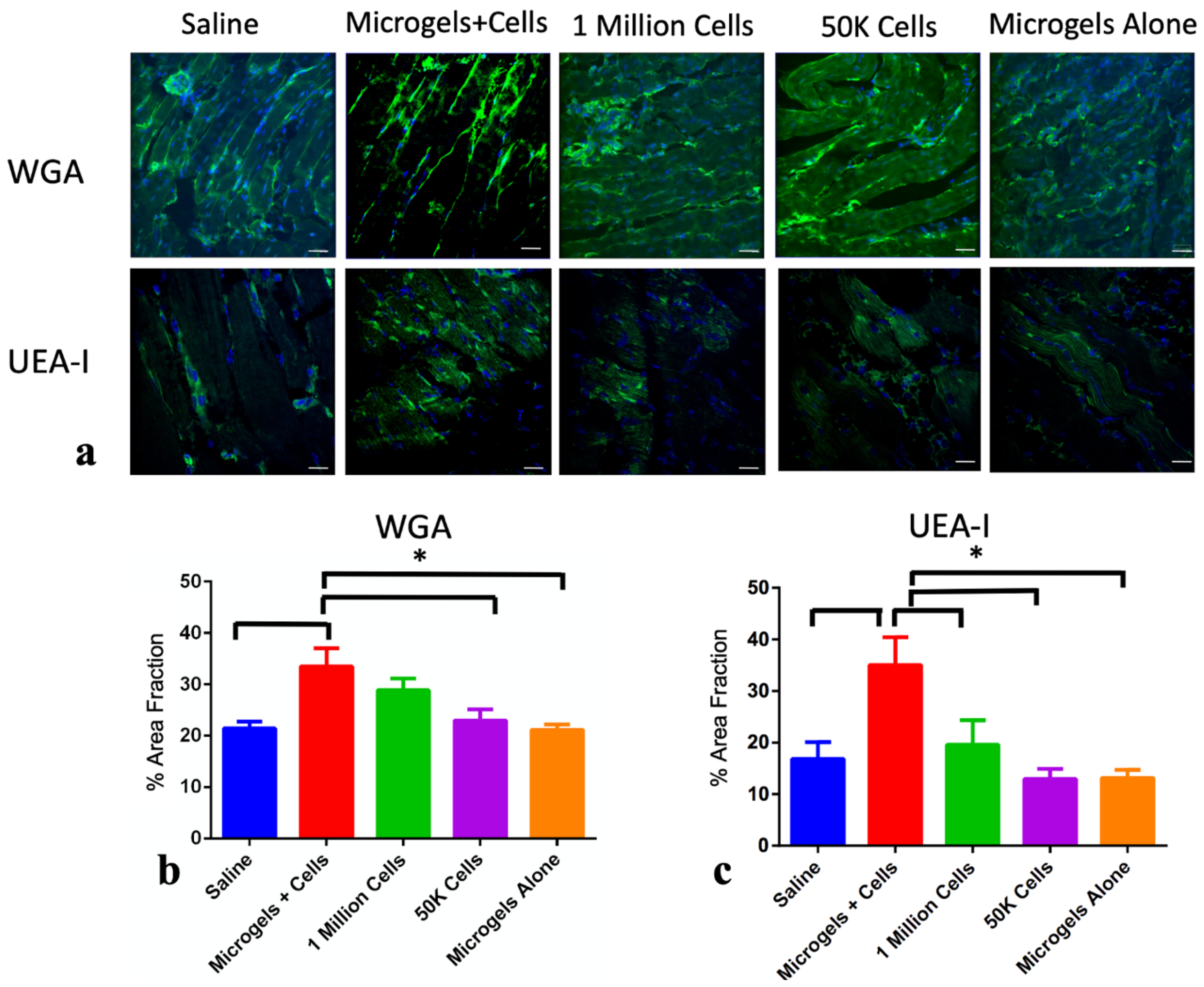


**Fig. S10**. Higher expression of WGA and UEA-I binding N-linked sugars in the treatment group. **(a)** Lectin immunohistochemical examination of N-linked sugars N-acetylglucosamine and a-linked fucoses with lectins WGA and UEA-I. Scale bar 50μm. **(b, c)** Percentage area fraction quantification based on the positive expression of the sugars in the upper gastrocnemius muscle tissue. Error bars represent SD $*p<0.05$ (One-way ANOVA).

**Table S1.** Integrins and extracellular matrix associated genes quantified using real-time PCR arrays at 24 hours.

| Refseq | Fold Change | 1mg | 2mg | 3mg | p value | | |
|---|---|---|---|---|---|---|---|
| NM_006988 | ADAMTS1 | -3.6619 | -4.7941 | -5.6452 | 0.00011 | 0.000028 | 0.000015 |
| NM_139025 | ADAMTS13 | -1.03 | -1.6833 | -2.1391 | 0.914939 | 0.187613 | 0.015388 |
| NM_080629 | COL11A1 | -2.9267 | -3.0271 | -2.8357 | 0.000435 | 0.002107 | 0.00037 |
| NM_004370 | COL12A1 | -3.0369 | -2.7984 | -5.6452 | 0.000002 | 0.000003 | 0.000001 |
| NM_021110 | COL14A1 | -2.3993 | -2.0064 | -2.2716 | 0.011728 | 0.016806 | 0.012408 |
| NM_001855 | COL15A1 | 2.8364 | -1.3237 | -1.4746 | 0.266421 | 0.014181 | 0.013105 |
| NM_001856 | COL16A1 | -1.4941 | -1.9336 | -2.6704 | 0.039006 | 0.014965 | 0.006407 |
| NM_000088 | COL1A1 | -2.3608 | -2.0344 | -2.8554 | 0.026649 | 0.035922 | 0.013371 |
| NM_001846 | COL4A2 | -2.6195 | -3.1556 | -4.3581 | 0.000401 | 0.000228 | 0.000127 |
| NM_000093 | COL5A1 | -2.7881 | -2.6474 | -4.1517 | 0.002718 | 0.002363 | 0.000565 |
| NM_001850 | COL8A1 | -1.8352 | -1.8208 | -2.23 | 0.005885 | 0.000295 | 0.000022 |
| NM_001901 | CTGF | -23.6855 | -80.5191 | -32.53 | 0.001326 | 0.001174 | 0.001262 |
| NM_001903 | CTNNA1 | -2.2751 | -2.058 | -1.8408 | 0.029556 | 0.05451 | 0.028189 |
| NM_001331 | CTNND1 | -2.0222 | -2.4873 | -2.8031 | 0.008589 | 0.005667 | 0.001418 |
| NM_002026 | FN1 | -2.6993 | -2.3208 | -3.8203 | 0.010269 | 0.013668 | 0.004137 |
| NM_000201 | ICAM1 | 14.1304 | 5.1292 | 11.7368 | 0 | 0.000016 | 0 |
| NM_181501 | ITGA1 | -3.9612 | -2.587 | -3.4992 | 0.000013 | 0.000014 | 0.000008 |
| NM_002203 | ITGA2 | 2.3306 | 4.0617 | 2.0368 | 0.053691 | 0.009823 | 0.004228 |
| NM_002204 | ITGA3 | -2.0457 | -2.3261 | -2.5263 | 0.001073 | 0.000115 | 0.000023 |
| NM_000885 | ITGA4 | -9.9585 | -5.4943 | -4.4806 | 0.004433 | 0.005348 | 0.005646 |
| NM_000210 | ITGA6 | -4.375 | -3.701 | -5.2916 | 0.000032 | 0.000036 | 0.000025 |
| NM_002206 | ITGA7 | -1.8999 | -2.111 | -2.5031 | 0.020846 | 0.015382 | 0.00258 |
| NM_002210 | ITGAV | -3.3853 | -2.8904 | -3.3105 | 0.005408 | 0.009179 | 0.00483 |
| NM_002211 | ITGB1 | -3.6619 | -2.4249 | -3.0392 | 0.00001 | 0.000001 | 0.000014 |
| NM_000211 | ITGB2 | -2.5478 | -2.375 | -2.538 | 0.004429 | 0.007681 | 0.003755 |
| NM_000212 | ITGB3 | 1.4182 | 2.5293 | 1.4773 | 0.051549 | 0.002152 | 0.006774 |
| NM_000216 | ANOS1 | -7.3748 | -3.701 | -4.9486 | 0.000168 | 0.000253 | 0.000187 |
| NM_005559 | LAMA1 | -6.2446 | -6.4588 | -5.0177 | 0.000031 | 0.00002 | 0.000001 |
| NM_000426 | LAMA2 | -3.3079 | -3.0623 | -4.7252 | 0.002153 | 0.000152 | 0.000072 |
| NM_000227 | LAMA3 | -6.6928 | -5.7408 | -2.4915 | 0.003105 | 0.012723 | 0.011063 |
| NM_002291 | LAMB1 | -2.8665 | -2.4193 | -2.8953 | 0.027368 | 0.040627 | 0.024394 |
| NM_000228 | LAMB3 | 1.7951 | 2.6064 | 1.598 | 0.028058 | 0.001556 | 0.000013 |
| NM_002293 | LAMC1 | -2.9813 | -2.9375 | -3.6059 | 0.001454 | 0.001681 | 0.000757 |
| NM_002421 | MMP1 | 1.7541 | 2.6004 | 1.1431 | 0.009561 | 0 | 0.476348 |
| NM_002425 | MMP10 | -1.5255 | -1.6755 | -2.0098 | 0.010072 | 0.170092 | 0.002702 |
| NM_005940 | MMP11 | 1.867 | 3.9233 | 2.4447 | 0.068653 | 0.022442 | 0.006527 |
| NM_002427 | MMP13 | 1.3232 | 3.3529 | 1.1944 | 0.321517 | 0.000702 | 0.552008 |
| NM_005941 | MMP16 | -3.7216 | -3.5176 | -2.9493 | 0.003553 | 0.003844 | 0.003728 |
| NM_002423 | MMP7 | 1.6595 | 3.1794 | 1.1944 | 0.125333 | 0.019935 | 0.552008 |
| NM_000615 | NCAM1 | -4.3649 | -1.1958 | -3.4911 | 0.000178 | 0.965691 | 0.007046 |
| NM_000442 | PECAM1 | 5.9686 | 6.5074 | 8.035 | 0 | 0.003749 | 0.000007 |
| NM_003119 | SPG7 | -1.7727 | -2.058 | -1.5337 | 0.030174 | 0.019686 | 0.021442 |
| NM_000358 | TGFBI | -2.3122 | -1.8124 | -2.4234 | 0.000312 | 0.000524 | 0.000622 |
| NM_003246 | THBS1 | -29.4311 | -94.0002 | -30.1419 | 0.000001 | 0 | 0.000001 |
| NM_003247 | THBS2 | -2.2438 | -1.9742 | -2.4403 | 0.002872 | 0.002641 | 0.000612 |
| NM_007112 | THBS3 | -1.4005 | -1.8293 | -2.298 | 0.000969 | 0.009572 | 0.00025 |
| NM_003255 | TIMP2 | -2.656 | -1.9879 | -2.4067 | 0.001778 | 0.008252 | 0.001295 |
| NM_000362 | TIMP3 | -1.8868 | 1.6534 | -2.7518 | 0.028095 | 0.071822 | 0.008229 |
| NM_002160 | TNC | -2.416 | -3.9393 | -3.3644 | 0.00289 | 0.000941 | 0.001179 |

| | | | | | | | |
|---|---|---|---|---|---|---|---|
| NM_001078 | VCAM1 | -1.1167 | -6.733 | -1.7577 | 0.018539 | 0 | 0.000067 |
| NM_004385 | VCAN | -4.5397 | -4.3307 | -9.3202 | 0.000103 | 0.000115 | 0.000095 |
| NM_000638 | VTN | -3.0864 | -1.3055 | -2.298 | 0.000318 | 0.015178 | 0.000148 |

**Table S2.** Integrins and extracellular matrix associated genes quantified using real-time PCR arrays at 96 hours.

| Refseq | Fold Change | 1mg | 2mg | 3mg | *p* value | | |
|---|---|---|---|---|---|---|---|
| NM_006988 | ADAMTS1 | 1.4313 | 2.0404 | -1.3827 | 0.291948 | 0.092411 | 0.261684 |
| NM_139025 | ADAMTS13 | -4.5714 | -2.5336 | -15.0062 | 0.006557 | 0.018146 | 0.002676 |
| NM_003278 | CLEC3B | -3.0794 | 2.6187 | -2.3416 | 0.491895 | 0.00584 | 0.043711 |
| NM_001843 | CNTN1 | -2.1033 | -2.1652 | -1.9108 | 0.224198 | 0.268665 | 0.263192 |
| NM_080629 | COL11A1 | -30.2593 | -6.11 | -40.6211 | 0.041776 | 0.031804 | 0.019703 |
| NM_004370 | COL12A1 | -1.6927 | -3.8491 | -32.9184 | 0.127129 | 0.028925 | 0.009252 |
| NM_021110 | COL14A1 | -35.736 | -68.9674 | -22.1232 | 0.016574 | 0.010692 | 0.004751 |
| NM_001855 | COL15A1 | -5.2029 | -3.8669 | -30.9276 | 0.012388 | 0.017439 | 0.006435 |
| NM_001856 | COL16A1 | -1.3622 | -2.0249 | -13.0939 | 0.287203 | 0.081803 | 0.015623 |
| NM_000088 | COL1A1 | -1.5397 | -2.0016 | -12.359 | 0.187439 | 0.105892 | 0.018435 |
| NM_001846 | COL4A2 | 1.031 | -1.6638 | -8.6188 | 0.878401 | 0.259828 | 0.019736 |
| NM_000093 | COL5A1 | -1.4267 | -2.2056 | -7.858 | 0.340562 | 0.195862 | 0.03415 |
| NM_001848 | COL6A1 | 2.2253 | 1.612 | -2.2153 | 0.079158 | 0.36556 | 0.158815 |
| NM_001849 | COL6A2 | 1.7022 | 1.4065 | -2.6223 | 0.281298 | 0.421352 | 0.152427 |
| NM_000094 | COL7A1 | 1.06 | -1.531 | -7.4858 | 0.846218 | 0.316742 | 0.007285 |
| NM_001850 | COL8A1 | 1.0454 | -1.5346 | -10.1318 | 0.849889 | 0.132168 | 0.003106 |
| NM_001901 | CTGF | -2.0839 | -2.1954 | -6.5318 | 0.059351 | 0.054333 | 0.012905 |
| NM_001331 | CTNND1 | -1.0541 | -1.4418 | -3.2735 | 0.877398 | 0.526991 | 0.002471 |
| NM_004425 | ECM1 | 1.7703 | 2.7362 | -2.0527 | 0.225514 | 0.246841 | 0.090306 |
| NM_002026 | FN1 | -1.4333 | -1.6871 | -7.5903 | 0.276179 | 0.187048 | 0.037213 |
| NM_000201 | ICAM1 | 2.0667 | 3.0784 | 1.3279 | 0.050834 | 0.067378 | 0.539919 |
| NM_181501 | ITGA1 | -2.8205 | -2.5336 | -9.4971 | 0.026033 | 0.03104 | 0.008707 |
| NM_002203 | ITGA2 | 2.9981 | 3.5707 | -1.0576 | 0.036151 | 0.02698 | 0.915943 |
| NM_002204 | ITGA3 | 1.0773 | 1.3586 | -2.6162 | 0.874485 | 0.292609 | 0.046838 |
| NM_000885 | ITGA4 | -1.2858 | -1.7305 | -6.3827 | 0.28309 | 0.096872 | 0.000191 |
| NM_002205 | ITGA5 | 1.6611 | 2.0383 | -1.6789 | 0.286055 | 0.024094 | 0.157791 |

| | | | | | | | |
|---|---|---|---|---|---|---|---|
| NM_000210 | ITGA6 | -1.0135 | -1.0603 | -3.35 | 0.974391 | 0.659283 | 0.012378 |
| NM_002206 | ITGA7 | -1.1116 | -1.5887 | -6.2513 | 0.912247 | 0.065609 | 0.004835 |
| NM_003638 | ITGA8 | -1.4201 | -1.7225 | -5.7924 | 0.244077 | 0.077203 | 0.009736 |
| NM_002210 | ITGAV | 1.436 | 2.4463 | -7.2308 | 0.526281 | 0.58429 | 0.014234 |
| NM_002211 | ITGB1 | -1.1969 | -1.1156 | -5.113 | 0.633716 | 0.758885 | 0.082413 |
| NM_000211 | ITGB2 | -1.0565 | 1.2076 | -3.8749 | 0.835744 | 0.647437 | 0.034589 |
| NM_000212 | ITGB3 | 1.3954 | 2.003 | -2.0574 | 0.459113 | 0.036211 | 0.060875 |
| NM_000213 | ITGB4 | 2.5859 | 1.0586 | -1.5628 | 0.224846 | 0.944684 | 0.303658 |
| NM_002213 | ITGB5 | 1.8474 | 2.21 | -1.2318 | 0.215782 | 0.013062 | 0.490664 |
| NM_000216 | ANOS1 | 1.284 | -1.7506 | -6.825 | 0.659121 | 0.221638 | 0.060856 |
| NM_005559 | LAMA1 | 2.4072 | 1.8906 | 1.194 | 0.363288 | 0.491004 | 0.855663 |
| NM_000426 | LAMA2 | -1.6051 | -2.1954 | -15.357 | 0.469286 | 0.164689 | 0.06037 |
| NM_000227 | LAMA3 | 3.0469 | 1.5788 | -2.0151 | 0.301858 | 0.175204 | 0.115915 |
| NM_002291 | LAMB1 | 1.1981 | -1.2522 | -6.0523 | 0.801917 | 0.434265 | 0.054972 |
| NM_000228 | LAMB3 | 3.2506 | 2.5707 | -1.4717 | 0.186198 | 0.15207 | 0.39836 |
| NM_002293 | LAMC1 | -1.62 | -1.4154 | -7.8944 | 0.106215 | 0.193932 | 0.009004 |
| NM_002421 | MMP1 | 9.3223 | 11.9491 | 9.0579 | 0.000272 | 0.012838 | 0.000667 |
| NM_002425 | MMP10 | 5.5175 | 3.4633 | 5.5245 | 0.000688 | 0.0464 | 0.124706 |
| NM_002426 | MMP12 | 7.2973 | 5.2011 | 8714.3504 | 0.37064 | 0.374811 | 0.373899 |
| NM_002427 | MMP13 | -2.6561 | -3.1264 | -1.5166 | 0.072851 | 0.061098 | 0.215509 |
| NM_005941 | MMP16 | -1.3311 | -1.3328 | -4.4203 | 0.442529 | 0.412449 | 0.059845 |
| NM_004530 | MMP2 | -1.0565 | -1.1234 | -2.5388 | 0.652923 | 0.504051 | 0.008733 |
| NM_002422 | MMP3 | 6.0658 | 4.7201 | 10.2142 | 0.059123 | 0.005099 | 0.10578 |
| NM_002424 | MMP8 | -1.8058 | -3.1627 | -2.4637 | 0.319593 | 0.061837 | 0.10789 |
| NM_000615 | NCAM1 | 1.8974 | 1.4098 | -2.2 | 0.07635 | 0.416929 | 0.050517 |
| NM_000442 | PECAM1 | 3.8478 | 5.6784 | 1.9442 | 0.022118 | 0.027027 | 0.058111 |
| NM_003919 | SGCE | -2.0648 | -1.4023 | -5.4673 | 0.175387 | 0.419069 | 0.038389 |
| NM_003118 | SPARC | -1.5648 | -1.531 | -4.2698 | 0.196233 | 0.409777 | 0.037024 |
| NM_000582 | SPP1 | 1.4413 | 2.9735 | 1.0371 | 0.111107 | 0.040731 | 0.786489 |

| | | | | | | | |
|---|---|---|---|---|---|---|---|
| NM_000358 | TGFBI | 1.8799 | 2.3546 | -1.581 | 0.486057 | 0.199921 | 0.257902 |
| NM_003246 | THBS1 | -2.9268 | -2.7661 | -10.5134 | 0.026908 | 0.027556 | 0.008417 |
| NM_003247 | THBS2 | -1.5903 | -2.3804 | -11.1128 | 0.185701 | 0.067659 | 0.010294 |
| NM_007112 | THBS3 | -1.583 | -3.055 | -11.1643 | 0.060015 | 0.000989 | 0.000307 |
| NM_003255 | TIMP2 | -1.0348 | 1.2853 | -3.162 | 0.767286 | 0.693697 | 0.107756 |
| NM_000362 | TIMP3 | -1.8142 | -1.1985 | -4.3797 | 0.24341 | 0.960682 | 0.03186 |
| NM_002160 | TNC | -1.4135 | -1.1929 | -8.6988 | 0.286224 | 0.898452 | 0.03333 |
| NM_001078 | VCAM1 | -1.9761 | -1.6072 | -10.7343 | 0.076188 | 0.300433 | 0.011799 |
| NM_004385 | VCAN | 1.3417 | -1.3115 | -9.9233 | 0.700811 | 0.823258 | 0.183297 |
| NM_000638 | VTN | -2.3663 | -2.5689 | -5.7924 | 0.222895 | 0.025737 | 0.009688 |
